# Quantification of nanocondensates formation at the single molecule level

**DOI:** 10.1101/2024.05.05.592604

**Authors:** Justin Houx, Thomas Copie, Yann Gambin, Emma Sierecki

## Abstract

Understanding the molecular mechanisms of biomolecular condensate formation through liquid-liquid phase separation is crucial for deciphering cellular cues in normal and pathological contexts. Recent studies have highlighted the existence of sub-micron assemblies, known as nanocondensates or mesoscopic clusters, in the organization of a significant portion of the proteome. However, as smaller condensates are invisible to classical microscopy, new tools must be developed to quantify their numbers and properties. Here, we establish a simple analysis framework using single molecule fluorescence spectroscopy to quantify the formation of nanocondensates diffusing in solution. We used the low-complexity domain of TAR DNA-binding protein 43 (TDP-43) as a model system to show that we can recapitulate the phase separation diagram of the protein in various conditions. Single molecule spectroscopy reveals rapid formation of TDP-43 nanoclusters at ten-fold lower concentrations than described previously by microscopy. We demonstrate how straightforward fingerprinting of individual nanocondensates provides an exquisite quantification of their formation, size, density, and their temporal evolution. Overall, this study highlights the potential of single molecule spectroscopy to investigate the formation of biomolecular condensates and liquid-liquid phase separation mechanisms in protein systems.

## Introduction

The compartmentalization of biomolecules is essential to their biological function in eukaryotic cells. Membraneless organelles and other biocondensates are increasingly recognized as important cellular environments, in addition to membrane-bounded organelles. They play an important role in the assembly of small nuclear ribonucleoprotein and ribosome biogenesis, genome organization or regulation of gene expression, among other cellular processes^1–3^. Biocondensates are formed through condensation of proteins, nucleic acids and other intracellular components, forming a dense liquid phase of biomolecules inside a dilute phase via a process of liquid-liquid phase separation (LLPS)^4,5^. Due to their liquid nature, biocondensates are often highly dynamic and their existence is very reactive to changes in their cellular environment.

Pure *in vitro* reconstitutions as well as high resolution cell imaging have already provided a wealth of information on how weak multivalent interactions between intracellular components contribute to the process of LLPS, for example^5–8^. Current techniques tend to generate information on microscopic-sized condensates, however the existence of sub-micron assemblies, referred to as nanocondensates, is increasingly recognized^9^. These assemblies exist at subsaturated physiological concentrations making them biologically relevant. Potentially, sub-micron organization could be widespread in the cell, with a recent study indicating that almost 20% of the proteome could be organized in nanocondensates^10^. Here, we used single molecule fluorescence spectroscopy to characterise the transition between homogenous solutions of monomeric proteins to condensates. This technique is particularly well-adapted to study heterogenous systems and provides the perfect tool to monitor the formation and evolution of nanocondensates^1^^1^.

First, phase transition was examined at the single molecule level with the low-complexity domain (LCD) of TAR DNA-binding protein 43 (TDP-43) as a model system, in the presence of a small chemical chaperone, trimethylamine *N-*oxide (TMAO), known to induce LLPS^12^. Single molecule fluorescence revealed the immediate formation of small condensates, at much lower protein concentrations that previously described in that system. We showed that our method enables precise mapping of LLPS and demonstrate reversibility upon 1,6-hexanediol addition or by dilution. We then explored the kinetics of condensation in detail and obtained an estimate of the physical sizes of the objects formed, validating that we are indeed observing nanocondensates in the 100 nm range. The use of a more physiological medium, a modified cell lysate, validated the trends observed with TMAO.

## Results

### Principle of single molecule detection of condensates formed through LLPS

LLPS is an inherently heterogenous process that requires the formation of mesoscopic assemblies that grow or coalesce on their way to form microscopic objects (Figure 1A). In the first steps of LLPS, we expect to observe the formation of local inhomogeneities of protein density, formation of very small nuclei that will recruit more proteins to grow. Tracking potentially rare and small inhomogeneities of concentrations in a high background of proteins, without immobilisation that could perturb those transitions, is difficult with fluorescence microscopy. However, single molecule spectroscopy methods were developed to measure protein diffusing in solution and have been used to study protein complexes and their growth^13,14^. In single molecule spectroscopy, fluorescent proteins freely diffusing in solution are detected as they travel through the small focal volume (typically 400nm wide, 1μm height (Figure 1B))^15^. As measurement focuses on this sub-micron volume, it is perfectly suited to detect small inhomogeneities in local concentrations. Typical fluorescence time traces are presented in Figure 1C. The number of photons captured by the photon counting detector is dependent on the number of labelled proteins present in the focal volume. Protein monomers in a homogenous solution appear as a flat baseline since the number of fluorescent proteins is constant throughout the solution (Figure 1C top). When a protein assembly crosses the focal volume, a sharp increase in fluorescence intensity is detected (Figure 1C bottom). Our group has previously characterized the oligomerization/aggregation of multiple proteins using this technique, including α-synuclein^16–18^. We developed a custom analysis to count the number of events and to report for each event, the peak intensity, the peak width (time taken to transit through the focal volume), and “total intensity” (calculated as the area under the curve) for each peak, or for the overall trace (Figure 1D)^16^. As we will demonstrate, these parameters contain interesting information about the condensates formed in LLPS.

**Fig. 1:**
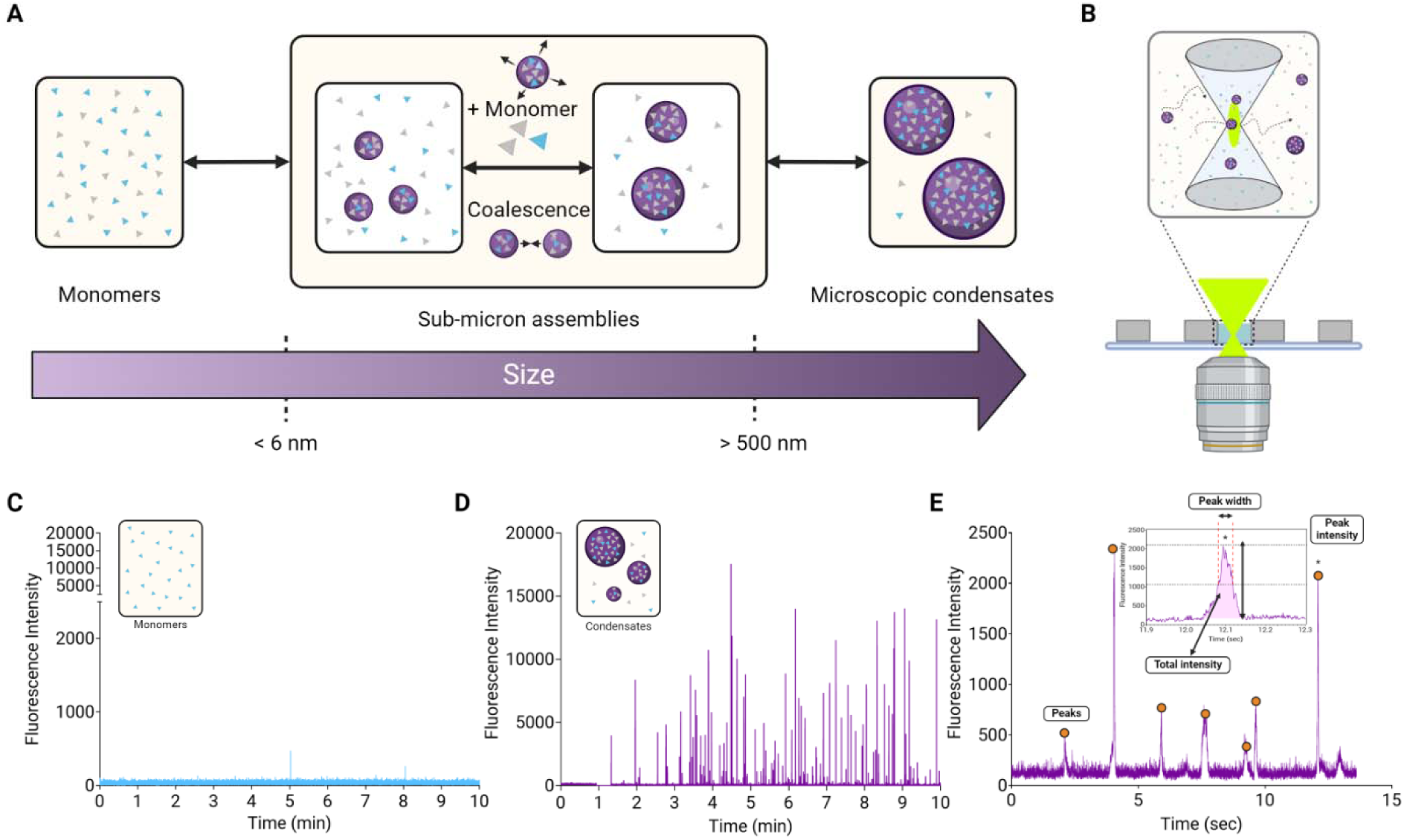
Single molecule approach to detect Liquid-Liquid Phase separation. A) Schematic of LLPS. TDP-43 LCD monomers are illustrated as triangles (blue, labeled with Atto647N and grey, unlabeled). Condensates are represented by purple spheres. From a monomeric solution, sub-micron assemblies are formed by nucleation and stabilized by growth/ coalescence, eventually leading to the formation of microscopic droplets. B) Schematic representation of the single molecule approach: a confocal volume, created by a 40x objective, is used to detect assemblies freely diffusing in a solution. C- D) Representative fluorescence traces of Atto647N-labeled TDP-43 LCD monomers (500 nM) in buffer without (C) and with 2M TMAO (D), measured over 10 minutes. The fluorescent peaks in D indicate the presence of TDP-43 LCD condensates. E) Example of graph showing a trace analysis which reports th total number of peaks (events) per measurement and (inset) the individual peak intensity, peak width (full width half-maximum), and total intensity (pink area under the curve). The inset shows a zoom in on the peak denoted by (*) in the trace.

### TDP-43 as a model system for LLPS

In this study, we used the low complexity domain (LCD) of the human TDP-43 as a model system (residues 263-414, Figure S1). TAR DNA-binding protein 43 (TDP-43) is normally a transcriptional repressor, also involved in splicing^19^. TDP-43 aggregates are commonly found in diseases such as amyotrophic lateral sclerosis, frontotemporal dementia, Alzheimer’s disease, and Limbic-predominant age-related TDP-43 encephalopathy (LATE)^19–22^. Recent studies have shown that the LCD of TDP-43 undergoes LLPS in a variety of conditions and strongly promotes the formation of amyloids^23^. In cells, TDP-43 LLPS is mediated by post-translational modifications, redox status, and presence of RNA while *in vitro* TDP-43 LCD was shown to form microscopic droplets upon variation of pH and salt concentration or in the presence of heparin, Zn^2+^ or ATP^24^. Here we chose to use a chemical chaperone, trimethylamine *N-*oxide (TMAO), to modulate TDP-43 LCD transition. Moosa et al. showed previously that TMAO effectively uncouples LLPS and aggregation, providing a good controlled system for our technique ^12,25^.

To extend our understanding, we also used a cell lysate as a surrogate physiological milieu and compared behaviour^26^. We used our cell-free protein expression system produced from the eukaryotic organism *Leishmania tarentolae*. Here this system is not used to produce proteins but rather to provide a more complex and more physiological environment, with realistic crowding. *Leishmania tarentolae* extracts (LTE) are orthogonal to human proteins and are presumed not to contain specific interactors for TDP-43; however, the extracts contain hundreds of different proteins, nucleic acids, lipids, and other components and therefore resemble cellular cytoplasm.

### Induction of LLPS of TDP-43 by Trimethylamine N-oxide (TMAO)

The chemical chaperone TMAO has a crowding effect on proteins and has been shown to stabilize the native fold of the protein both *in vivo* and *in vitro*^27^. In the case of TDP-43 LCD and other intrinsically disordered proteins, protein compaction has been observed upon titrating TMAO^12,25^. As a proof of concept, we used previously published experimental conditions to monitor LLPS as a function of TMAO concentration using single molecule spectroscopy^12^. Briefly, a fraction of TDP-43 LCD was labelled with the fluorophore Atto647N for fluorescence detection while the rest was kept untagged. The principle of these measurements is as follows: labelled TDP-43 LCD was diluted to low concentration (500 nM) and a measurement was started. After ∼60 sec, the detection was blocked and unlabelled TDP-43 LCD was rapidly added to the solution to reach the desired concentration of 1000 nM. The detection was then switched back on, allowing the measurement to continue. In the absence of TMAO, very few peaks were detected (Figure 1B); however, in the presence of 2M TMAO, the increase in protein concentration immediately resulted in peak detection within seconds (Figure 2A). 10-minute measurements were performed in triplicate with consistent results between the repeats (Figure S2). Analysis of the whole traces revealed significant changes in the number of events (Figure 2B) and total intensity (Figure 2C) depending on the concentration of TMAO. Both the number of peaks and total fluorescence intensity increased as a function of TMAO concentration. In contrast, the mean peak width and mean peak intensity did not vary significantly (Figure S3). These findings suggest that TMAO enhances LLPS of TDP-43 LCD leading to formation of a higher number of condensates within the first 10 minutes following exposure to TMAO.

**Fig. 2:**
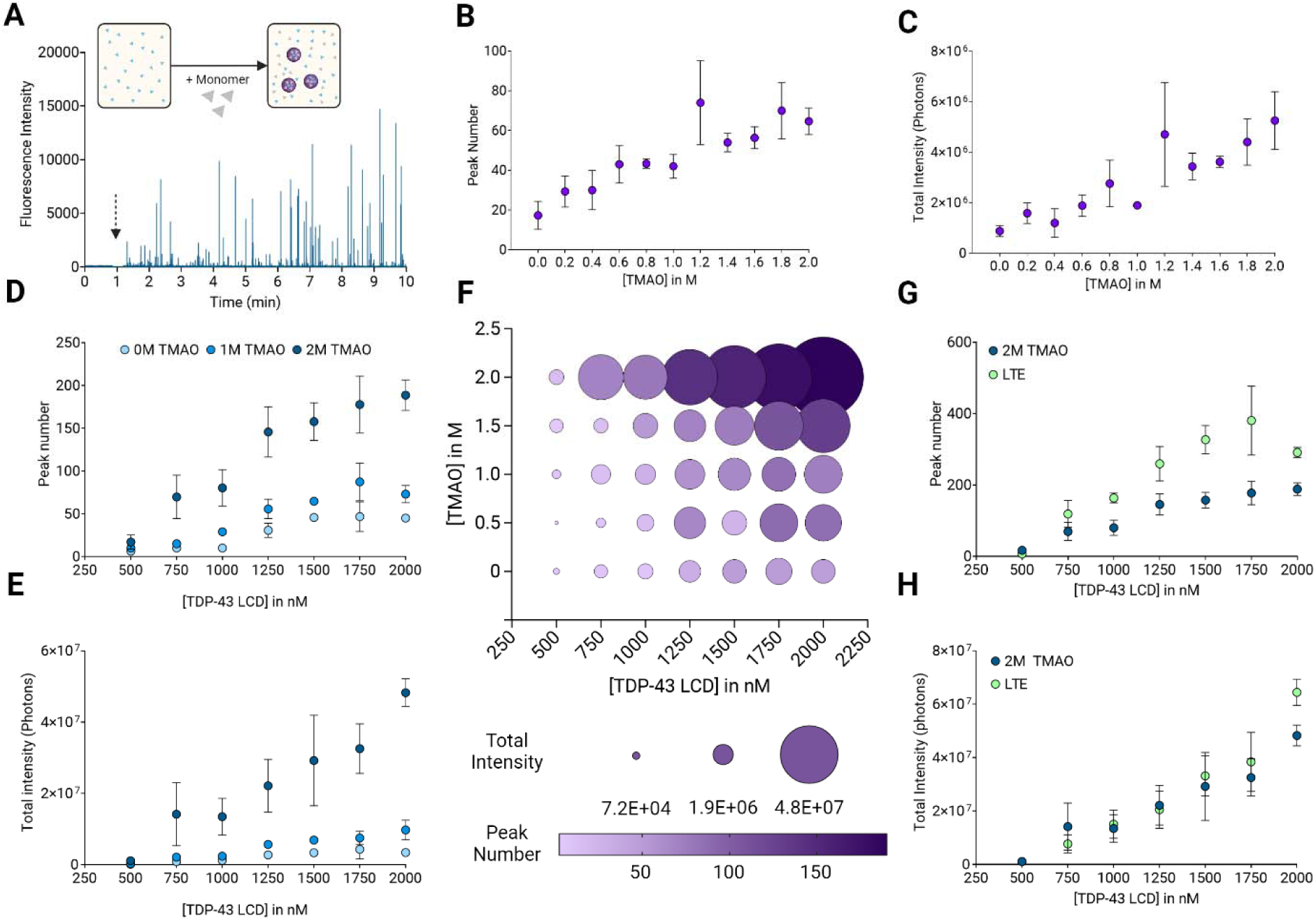
Effect of TMAO and TPD-43 LCD concentration on LLPS. A) Typical trace obtained using the experimental workflow. Initially, the sample solely contains Atto647N-labeled TDP-43 LCD (500 nM) in αβγ buffer. Immediate detection of nanocondensates occurs upon addition of unlabeled TDP-43 LCD (500 nM), indicated with the arrow. Note that during this period, the detection is blocked off. B) Number of peaks in 10-minute traces as a function of TMAO concentration. C) Total intensity in 10-minute traces as a function of TMAO concentration. D) Number of peaks in 10-minute traces as a function of TDP-43 LCD concentration in buffer containing 0, 1, or 2M TMAO. E) Total intensity n 10-minute traces as a function of TDP-43 LCD concentration in buffer containing 0, 1, or 2M TMAO. F) Number of peaks (color coded by light to dark purple) and total intensity (size coded by size of circle) in traces upon varying concentrations of TMAO and TDP-43 LCD in buffer. G) Comparison of the number of peaks in 2M TMAO in αβγ buffer (dark blue) and lysate extract (CFL, green) at different concentrations of TDP-43 LCD H) Comparison of the total intensity of traces in 2M TMAO in αβγ buffer (dark blue) and lysate extract (CFL, green) at different concentrations of TDP-43 LCD. All data presented as mean ± SEM, for the conditions 0, 1, and 2M TMAO and CFL triplicate data is shown.

### LLPS of TDP-43 LCD as a function of TMAO and protein concentration

Protein concentration is always determinant in phase transition. Serial dilutions of unlabelled TDP-43 LCD were used to vary the final concentration of protein, in combination with different concentrations of TMAO. For each combination, 10-minute measurements were obtained and analysed as before. As expected, the number of events (Figure 2D) and total intensity (Figure 2E) within these events increased with higher protein concentration. This is particularly striking in the presence of 2M TMAO but was observed for each concentration of TMAO. This data can be summarized in a phase diagram presented in Figure 2F. Here the number of events and fluorescence intensity within these events are represented as a function of both TDP-43 and TMAO concentration. Not surprisingly, the single molecule phase diagram matches previous observations but reveals transition at lower concentrations than described for formation of macroscopic droplets (for example, Choi et al.^12^ used 20 µM of TDP-43).

The experiments were repeated using the cell lysate LTE as a buffer. Similar experiments were conducted with increasing concentrations of unlabelled TDP-43 added to labelled TDP-43 in LTE. We observed very similar behaviour compared to buffer with 2M TMAO, with very fast formation of assembly upon mixing (Figure S4). Although more events were detected in LTE compared to 2M TMAO (Figure 2G), the total intensity within the events vary similarly in LTE and 2M TMAO as a function of TDP43 concentration (Figure 2H). These remarkably similar findings suggest that the fraction of protein undergoing condensation is an intrinsic property of the protein, irrespective of the buffer. Further analysis reveals that the average peak widths were also almost identical in both conditions (Figure S5). There was however a clear difference in the intensity of the objects formed with the LTE assemblies being less bright (mean intensity of 3240± 316 vs. 1547± 36 photons per 10ms, for 2M TMAO and LTE respectively) (Figure S5 and S6). Assemblies in 2M TMAO also reach a higher maximal intensity (Figure S6).

### Reversibility of TDP-43 LCD LLPS

Reversibility of assembly formation is an important criterion to assess the liquid-like property of protein condensates. Two sets of experiments were performed to demonstrate the dissolution of the observed assemblies. First, 1,6-hexanediol was used to illustrate the reversible properties of LLPS. This aliphatic alcohol inhibits weak hydrophobic interactions and is widely used to dissolve droplets^5^. A reduction in number of events was observed as a function of the percentage of 1,6-hexanediol (Figure 3A), validating that hydrophobic interactions play an important role in the process. Surprisingly, the mean peak intensity slightly increased as a function of 1,6-hexanediol concentration (Figure 3B), mainly due to a loss of the objects with the lowest intensity (Figure S7). Meanwhile, the mean peak width decreased non significantly (Figure 3C). This suggests that the “smaller” events disappear readily while the brighter objects persist at higher 1,6-hexanediol concentration.

**Fig. 3:**
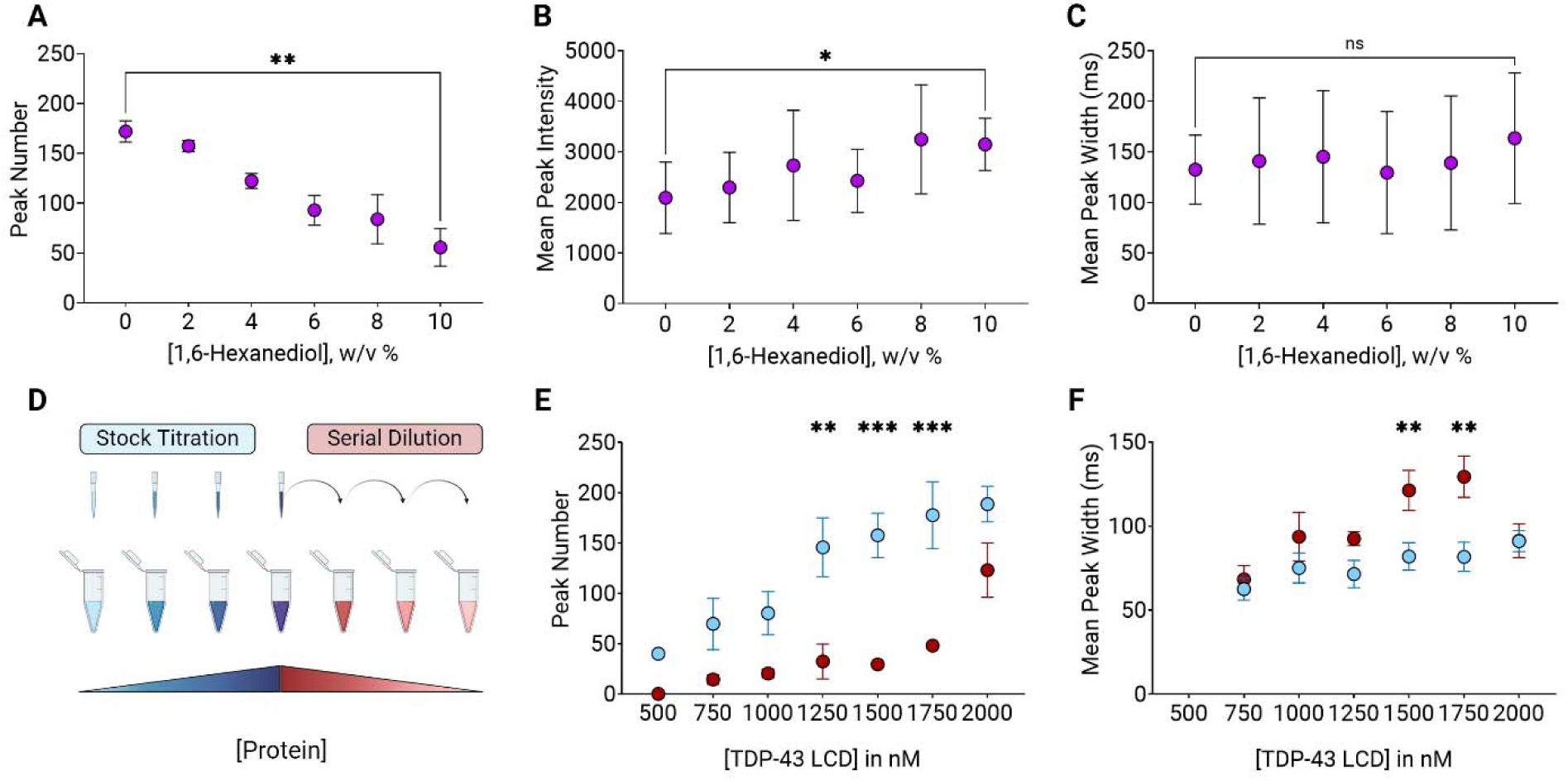
Reversibility of LLPS upon addition of 1,6-hexandiol and serial dilutions. A) Number of peaks in 10-minute traces for different concentrations of 1,6-hexanediol in 2M TMAO in αβγ buffer. B) Mean peak intensity for different concentrations of 1,6-hexanediol in 2M TMAO in αβγ buffer. C) Mean peak width for different concentrations of 1,6-hexanediol in 2M TMAO in αβγ buffer. D) schematic overview of the experimental layout for titrations (blue) and serial dilutions (red). For titrations, a starting solution at 20 µM in dd H_2_O is diluted in 2M TMAO in in αβγ buffer to the desired concentration. For dilutions, a solution containing 500nM labelled and 1500 nM unlabeled TDP-43 LCD is prepared and then serially diluted in a solution containing 500 nM labelled TDP-43 LCD in 2M TMAO, αβγ buffer. E) Number of peaks in 10-minute traces as a function of TDP-43 LCD concentration either upon titration (blue) or serial dilution (red). F) Mean peak width for different concentrations of TDP-43 LCD either upon titration (blue) or serial dilution (red). All data presented as mean ± SEM; statistics used Friedman test, p ≤ 0.05 (*), p ≤ 0.01 (**), p ≤ 0.001 (***), p ≤ 0.0001 (****), nonsignificant (n.s.).

Next, a dilution of a sample containing preformed assemblies was performed in buffer while keeping the concentration of labelled protein constant. Therefore, only the concentration of unlabelled protein was decreased (Figure 3D). Results showed that upon the dilution, the number of peaks significantly decreases (Figure 3E, red). Overlaying the data obtained for the direct titration and dilution experiments allows us to compare the two behaviours at identical protein concentrations. The number of peaks for the titration and the dilution at the lowest and highest concentration of protein were the same, however, the intermediate measurements tell a different story (Figure 3E). In the dilution experiments, the peak number instantly dropped from 100 to 50 for protein concentrations of 2000 and 1750 nM respectively (Figure 3E, red) while the number of peaks plateaued during the titration experiment (Figure 3E, blue). Further, the width of the peaks for the titration remained relatively constant for all concentrations of protein (Figure 3F, blue). In contrast, during dilution, the mean width of the peaks increased after the first dilution then gradually decreased until it reached similar values compared to the titration samples (Figure 3F, red). For most of the range of concentration explored, the width is significantly higher for samples obtained by dilution compared to those from titration. This again points to a difference of behaviour, either kinetic or thermodynamic, between smaller and larger objects, with the larger assemblies being more difficult to dissolve. Note that experiments with similar dilutions were carried out in LTE, the titration and dilution curves perfectly overlapped, and the observed hysteresis did not occur (Figure S8). This may be due to the fact that the brighter and wider objects that persisted in the presence of TMAO failed to form in LTE (Figure S6).

### Kinetics of TDP-43 LCD LLPS induction

Confident that we were able to observe the formation of condensates of TDP-43 LCD, we set out to examine the kinetics of the process, another poorly understood aspect of LLPS. Current research often uses long incubation times ranging from hours to multiple days^12,28^. This allows the droplets to grow enough to be observed using imaging techniques^29–33^. In contrast, our single molecule approach focusses on the early stages of phase transition. The data generated using 1000nM TDP-43 (total concentration) at varying TMAO concentrations (Figure 2) were analysed kinetically to determine when condensates form and start to grow in size rather than in number. To visualise this, a heat map was generated that indicates the number of events per minute as a function of TMAO concentration (Figure 4A). This analysis is inspired by the kymograph representations used in single molecule TIRF (Total Internal Reflection Fluorescence) microscopy experiments. It clearly shows that the time to generate a given concentration of objects (e.g. >10 events per time frame, orange colour) decreases as a function of TMAO concentration. The heat map further revealed that the concentration of objects varies over time, especially at low TMAO concentrations. This can also be seen on Figure 4B, which shows the number of events per minute in 4 experimental conditions. The presence of 2M TMAO or LTE led to the formation of a higher number of condensates at all time points.

**Fig. 4:**
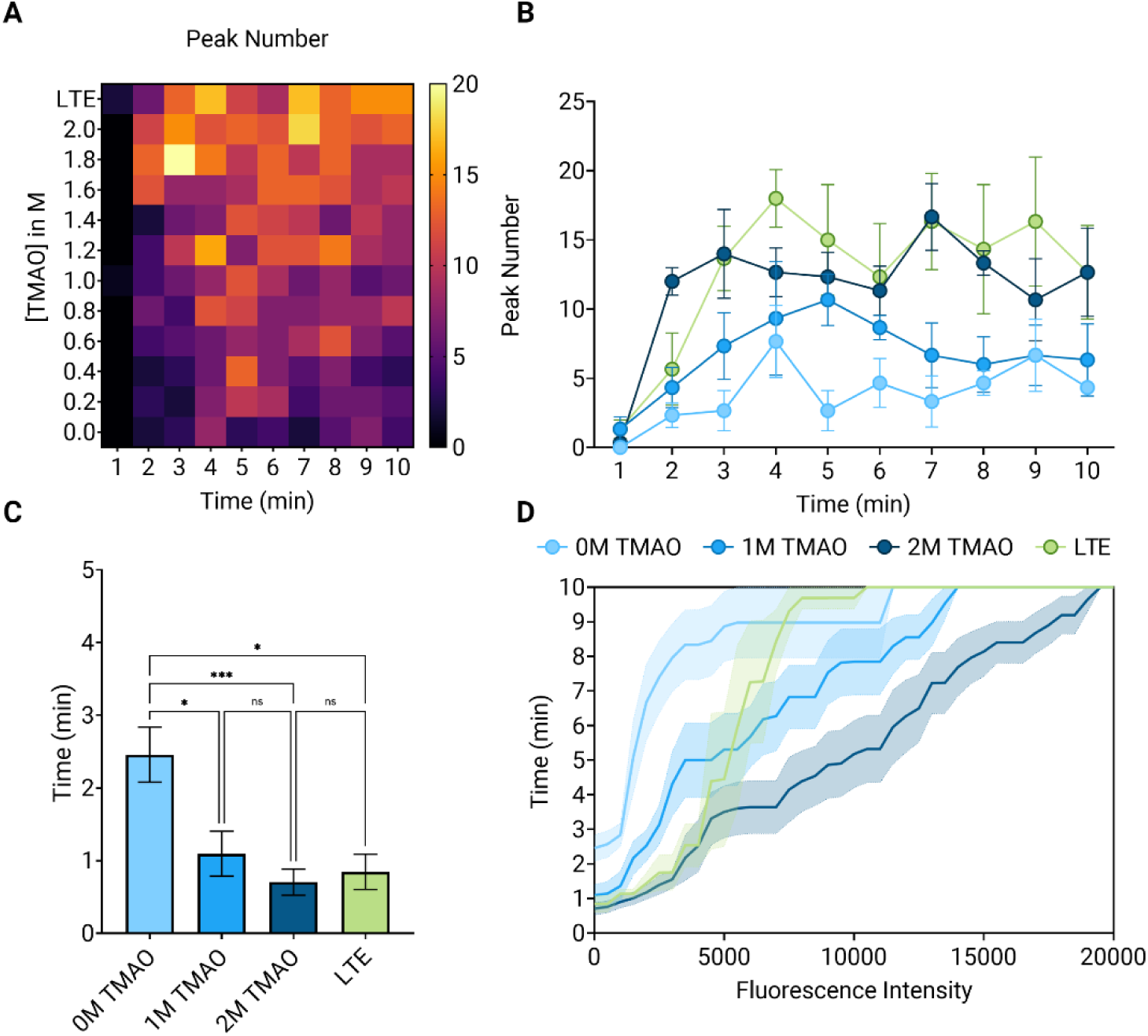
Kinetics of initial condensate formation. A) Heat map showing the number of events per minute for cell lysate (CFL) and a range of TMAO concentrations. Low to high number of peaks are indicated by a colour change from black/purple to orange/yellow. B) Number of peaks per minute in a 10-minute trace in four different experimental conditions (CFL in green, 0M TMAO in blue, 1M TMAO in purple, 2M TMAO in pink). C) Bar graph showing the time to the first peak detected in the trace under the same four experimental conditions as above (CFL in green, 0M TMAO in blue, 1M TMAO in purple, 2M TMAO in pink). D) Time to detect the first peak of a specific intensity as a function of fluorescence intensity, with shaded error area. The same experimental conditions are shown (CFL in green, 0M TMAO in blue, 1M TMAO in purple, 2M TMAO in pink). In all graphs the error bars are standard error of the mean (SEM) and statistics used Kruskal-Wallis test, p ≤ 0.05 (*), p ≤ 0.01 (**), p ≤ 0.001 (***), p ≤ 0.0001 (****), nonsignificant (n.s.).

We also asked whether TMAO affected the time it took to generate the first condensate (Figure 4C). The time to the first peak detected in a measurement was plotted as a function of the concentration of TMAO (Figure 4C, D). The first peak in the trace appeared after ∼60s at 1M TMAO compared to ∼40s at 2M TMAO (Figure 4C). This slight decrease indicates that TMAO has a limited effect on the kinetics of transition. Further, the positions for the first peak above a given intensity (Figure S9) were also calculated and plotted in Figure 4D. The delay in detecting the first peak of a given intensity increased as a function of intensity at all TMAO concentrations but was strongly reduced at higher TMAO concentration (Figure 4D). Note that the data overlapped once again for LTE and 2M TMAO for events below 4000 photons/ms but diverged afterwards. This is in line with the fact that LTE prevents the formation of highly fluorescent events (Figure S6). Overall, this suggests that TMAO plays a role in increasing the stability of the nuclei, allowing brighter objects to form faster.

### Evolution of the condensates fingerprints as a function of time

Next, we were interested in distinguishing different subgroups of condensates based on their size and fluorescence intensity. To extract this information, the intensity of each event was plotted against its peak width to fingerprint the properties of the condensates (Figure 5A). To compare these properties between different measurements, the events were categorized into four subgroups, depending on their position in different quadrants of the graph (Figure 5A). Thresholds for both the width and intensity were determined based on data obtained with the lowest TMAO (0M) and TDP-43 LCD (250nM unlabelled) concentrations with the requirement that 90% of the peaks have a peak width and intensity below the defined thresholds. The four quadrants resulting from these thresholds represent characteristic properties of peaks (Figure 5B). The bottom left quadrant (BL) indicates peaks with a small width and a low intensity. The top left quadrant (TL) corresponds to events with small widths and high intensity values. The top right quadrant (TR) has peaks with high values for both, and the bottom right quadrant (BR) indicates peaks with large widths but low intensities. The quadrant analysis was used to determine the number of peaks and percentage of peaks in each quadrant.

**Fig. 5:**
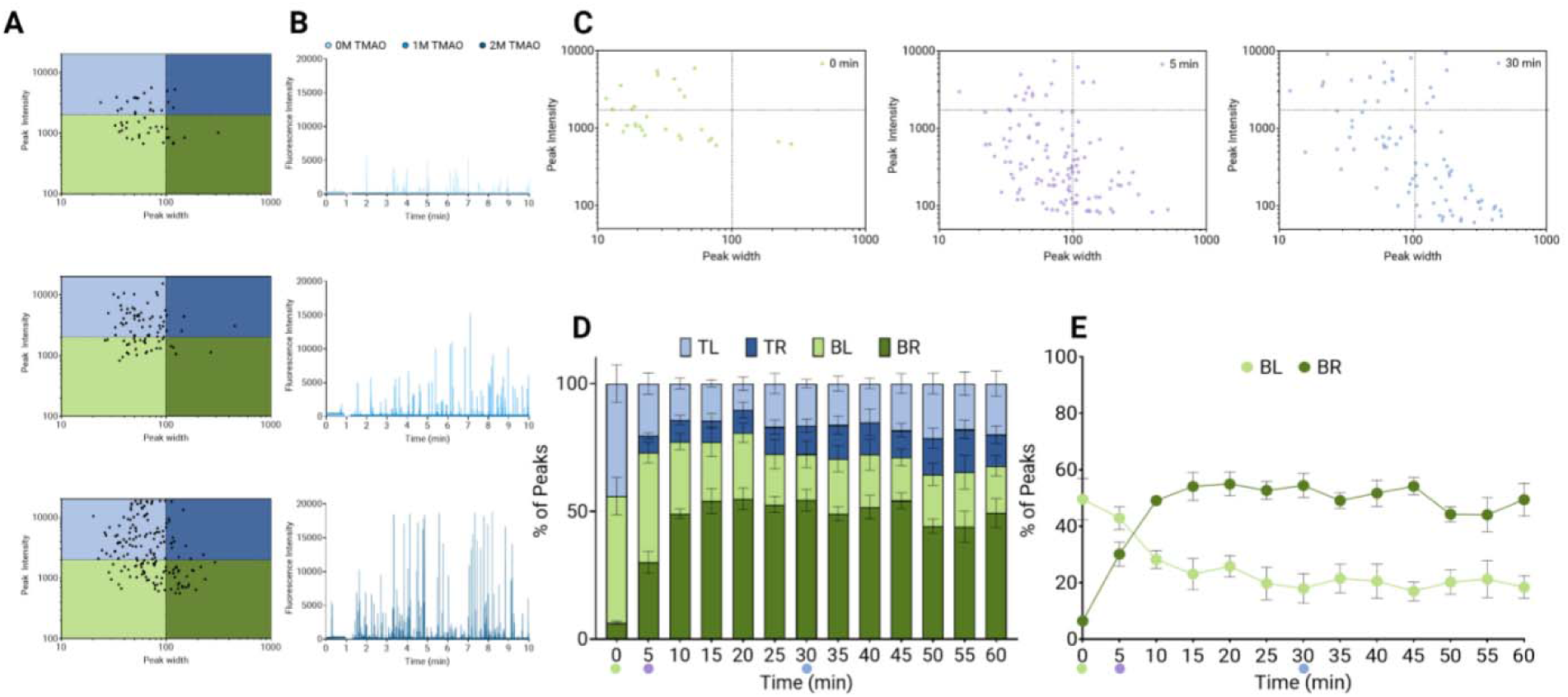
Fingerprinting the properties of the TDP-43 LCD condensates. A) Proof of principle of the quadrant analysis: scatter plots are generated showing the peak intensity as a function of peak width for each peak in a 10 -minute trace for 500 nM Atto674N-labeled TDP-43 LCD and 1500 nM unlabeled TDP-43 LCD in 0M (light blue), 1M (blue), and 2M (dark blue) TMAO in αβγ buffer. We defined 4 quadrants to distinguish between events. Four quadrants are defined: TL (light blue, > 2000 photons, < 100 ms), TR (dark blue, > 2000 photons, > 100 ms), BL (light green, < 2000 photons, < 100 ms), and BR (dark green, <2000 photons, > 100 ms). B) Corresponding raw fluorescence traces in 0M (light blue), 1M (blue), and 2M (dark blue) TMAO 1in αβγ buffer. C) Scatter plots showing the peak intensity as a function of peak width for 5-minute traces at different time points (0 min = green, 5 min = purple, and 30 min = blue). The vertical line at x = 100 and a horizontal line at y = 2000 are thresholds that define the different quadrants. D) Bar graph showing the percentage of peaks in each quadrant at different time points for 500 nM Atto674N-labeled and unlabeled TDP-43 LCD in 2M TMAO in αβγ buffer. Note that data at time 0 are obtained from 10-minute traces of 500 nM Atto674N-labeled TDP-43 LCD in 2M TMAO, αβγ buffer which represents the state before addition of unlabeled protein. E) Graph showing the transition between BL and BR over time. The percentage of peaks for BL (light green) and BR (dark green) in 5-minute traces are shown over time. In all graphs the mean is shown, and the error bars are standard error of the mean (SEM).

As a proof of concept, we compared the data acquired in the presence of 0, 1 and 2M TMAO. Representative scatter plots show a shift to other quadrants (Figure 5A) when compared to a sample in absence of TMAO. At 1M TMAO, the TL quadrant becomes populated and at 2M TMAO a higher percentage of events appear in the BR quadrant. Increasing the concentration of protein also influences the fingerprint of the events (Figure S10).

We then asked whether this analysis could give new insights on the dynamics of condensate formation. To do this, 1h time traces were acquired for 1000nM TDP-43 LCD (total) in the presence of 2M TMAO. The percentage of peaks in each quadrant was determined every 5 minutes (Figure 5 C-D). Initially, the majority of events have a fingerprint corresponding to the BL or BR quadrants (>70 %) with most events being fast diffusing (BL) (Figure 5D). Over time, the proportion of the top quadrants (fluorescently intense peaks) gradually increases from ∼20 to 35% (Figure 5D). The most dramatic change however is the rapid “conversion” between the BR and BL quadrant (Figure 5E), leading to a stable inversion of the proportion within 15 minutes. In the more physiological environment provided by the cell free extracts (LTE), the trends were similar, but the kinetics were different (Figure S11). The conversion of peaks in the BL quadrant to the BR quadrant occurs after a lag time of ∼ 30 minutes. In this case, the majority of events remain in the BL quadrant. Overall, the fingerprint of the objects changes over time, with faster transitions being observed in buffer/ TMAO conditions. The conversion of events from the BL to BR quadrant reflects an increase in the width rather than the intensity of the peaks, which suggests a change in the physical size of the objects.

### Correlation between signal width and condensate size

Intuitively, it makes sense that a larger object would take more time to cross the volume of detection. However, because of Brownian motion, the relationship between the two parameters is non-trivial. To be able to extract size information from our measurements, we first turned to mathematical simulations to examine how the signal changes as a function of the size of the object. For this, we simulated the random Brownian motion of a fluorescent sphere and modelled the confocal volume as a Gaussian function in the middle of box (Figure 6A, see Supplementary Information for details). The simulated signal resembles the experimental data, and we observe a smooth peak when a sphere “crosses” the virtual confocal volume (Figure 6B). Varying the size of the sphere, we plotted the peak width as a function of diameter; a linear relationship was observed, up to 400nm (Figure 6C, orange). Confident that there was a simple correlation between the size of an object and its time of residence in the confocal volume, we then set out to experimentally calibrate the experiment using a variety of homogeneous fluorescent objects of known diameter. Commercial fluorescent beads of 430, 200 and 100nm were used as well as liposomes (120nm) and purified Hepatitis B capsids (45nm). Multiple 5-minute fluorescence traces were acquired for all objects and the data was analysed as previously described. Figure 6C (blue) shows that the width of the individual peak increased as a function of size, and the average peak width is linearly proportional to the diameter of the object, in agreement with the simulations (see Supplementary Information and Figure S12). Based on this, we first conclude that almost all the objects we observed during TDP-43 LLPS are <400nm diameter (no width > 200ms). In our standard experiment using 1000nM TDP-43, the average size of the condensates is approximately 150nm based on our calibration curve. This is in agreement with the expected size of the nanocondensates, from tens to a few hundred nm^34^. We then estimated the average size of the condensates for different TMAO and TDP-43 concentrations (Figure S13). Strikingly, increase in TMAO concentration did not affect the physical size of the condensates (Figure 6D). However, higher protein concentration resulted in the formation of larger condensates in the presence of 2M TMAO or LTE (Figure 6E). By combining intensity and size information, we can estimate the protein concentration inside the nanocondensates (Figure 6F). The data demonstrates that the concentration in the dense phase does not vary for 2M TMAO when protein concentration increases, as expected for LLPS. This also suggests that at lower TMAO concentrations, we observe the formation of nuclei with different properties.

**Fig. 6:**
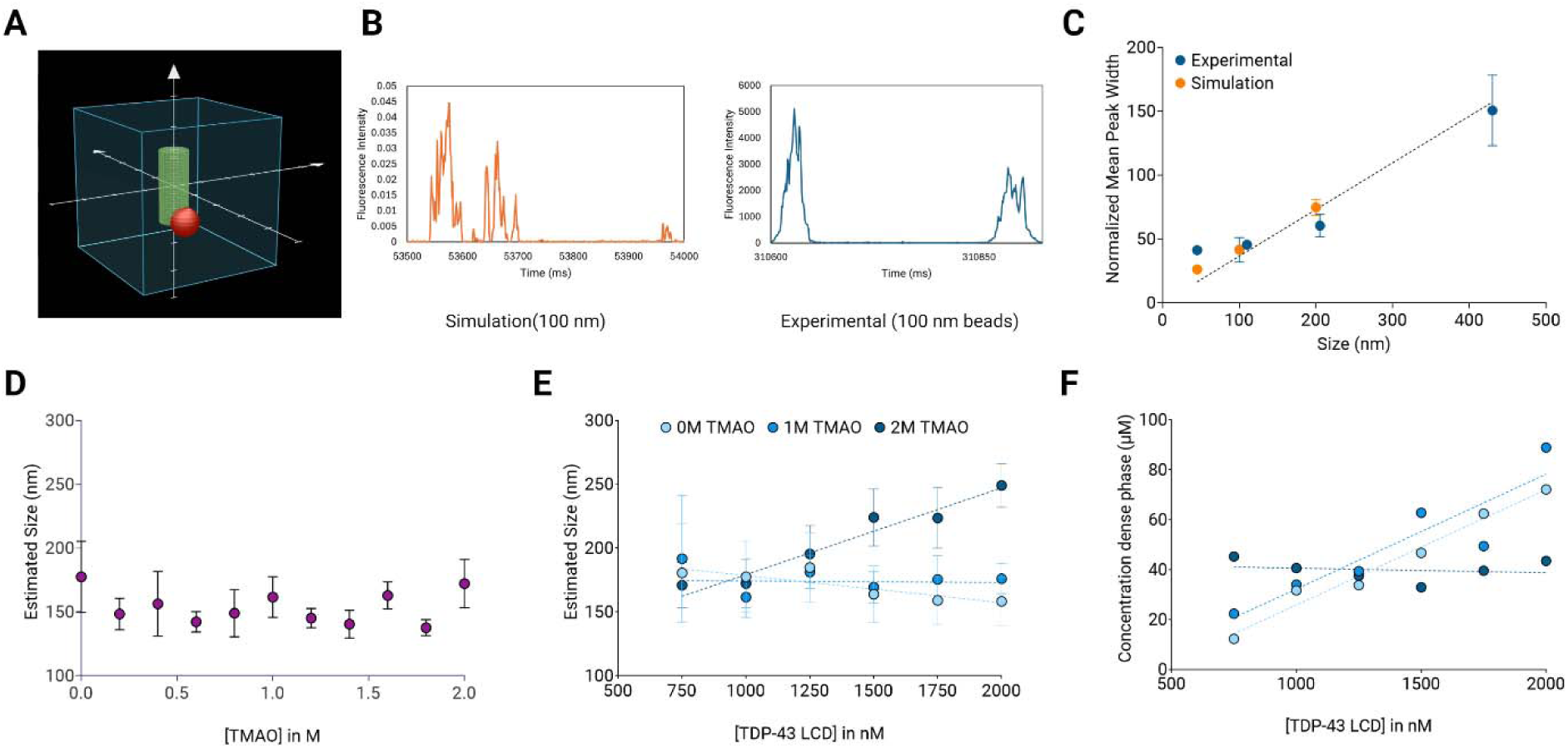
Calculation of condensate size and concentration of protein inside the dense phase. A) Time frame during the simulation. The cubic box has dimensions of 4 µm (blue), the gaussian confocal detection volume is shown as a cylinder (green), and the sphere is represented in red. B) Representative fluorescence traces showing peaks observed in simulations (orange) or experimentally (blue) for 100 nm beads. C) Calibration curve for size: the measured mean peak width values obtained experimentally (blue) or simulated (orange) are plotted as a function of the physical diameter of the object. A linear fit wa obtained after excluding the data points corresponding to 45 nm beads. Note that the peak width values for the simulations were normalized here to the experimental width observed for 100 nm beads. D) Estimated size of the condensates as a function of TMAO concentration, based on the previousl determined calibration curve. The data shown is the mean of three measurements and error bars indicate the standard error of the mean (SEM). E) Estimated size of the condensates as a function of TDP-43 LCD titration concentrations at three concentrations of TMAO (0, 1 and 2 M). Values are calculated from the mean peak width according to the previously determined calibration curve. The data shown is the mean of three measurements and error bars indicate the standard error of the mean (SEM). F) Estimated concentration of protein inside the condensates as a function of “bulk” TDP-43 LCD concentration, at three concentrations of TMAO (0,1 and 2M).

## Discussion

In this study, we investigated the initial formation of TDP-43 LCD condensates through liquid-liquid phase separation (LLPS) using single molecule spectroscopy. Our novel method detects the formation of sub-micron sized TDP-43 condensates. We could describe TDP-43 LCD phase separation at the single molecule level, at low concentrations of protein, and within the first minutes of nanocondensate formation. Indeed, we could characterize the condensates formed at concentrations ranging from 500 to 2000 nM, emphasizing the sensitivity of the method. LLPS of TDP-43 LCD could be induced by TMAO in a concentration dependent manner and the effects of both protein and TMAO concentrations could be summarized in a simplified phase diagram. Further, we showed that the protein also phase-separates in a crowded cell lysate, mimicking the cell environment. Interestingly, we noted that some properties of the condensates were insensitive to the milieu, while others (such as the maximal intensity) varied between “pure” buffer and cell lysate. Most of the nanocondensates were dissolved either by dilution or by addition of 1,6-hexanediol in a perfectly reversible manner when a more physiological buffer (LTE) was used. This method has the potential to become a simple tool that investigates the formation of biomolecular condensates as well as the mechanisms of LLPS under physiological conditions.

Previous studies have examined the LLPS of TDP-43 at concentrations varying from 15-600µM^12,28,33,35^; Garg and Bhat^28^ determined the saturation concentration to be 6.6µM at pH 7.4, in the presence of NaCl ^28^. Our study examined the behaviour of the protein at a 10-fold lower concentration, well within the sub-saturated regime, where nanocondensation is thought to occur. Indeed, protein nanocondensation is increasingly described, with recent examples being reported in the literature for FUS, c-GAS or α-synuclein^36–38^. Here we show that nanocondensates exist far below the concentration where microscopic condensation is detected (Figure 2). The average size of the objects in this study ranges from 100 to 250nm for protein concentration up to 2µM (Figure 7C-D). This agrees with the expected sizes of nanocondensates, that measure tens to hundreds nm in diameter. Recently, Amico et al.^39^ reported that FUS nanocondensates formed *in vitro* measured between 80 and 130nm, for the same concentrations range as our study. Nanoclusters of α-synuclein were also characterized and found to measure around 70nm in diameter^38^. Interestingly, in our case the size of the condensates was independent of TMAO concentration, and proportional to the concentration of protein only, in 2M TMAO and LTE (Figure 6, Figure S14).

The study of nanocondensation is gaining interest due to its biological relevance, but also because it is not well described by the Flory-Huggins model which is often used to explain the liquid-liquid phase separation of protein^9^. This theoretical framework predicts that in sub-saturated solutions, clusters of > 3-5 proteins should not exist. We show that at early time points (<10 min), the number of condensates formed increased as a function of protein concentration. The fraction of protein in these condensates appears invariant, as the total intensity in the condensates varies linearly with the bulk concentration (Figure 2). This holds true at all TMAO concentrations. Fitting the data to a linear fit yields a measure of the fraction of protein in the condensates and this volume fraction depends on the concentration of TMAO; the more TMAO, the more proteins are sequestered in the dense phase (Figure S15). Interestingly, there is a remarkable overlap of data between LTE and 2M TMAO when comparing the total intensity in the peaks (or volume fraction) (Figure 2H). Yet, the characteristics of the condensates formed are strikingly different between the two conditions. The objects detected in 2M TMAO are significantly brighter and smaller compared to the ones formed in LTE (Figure S6). This indicates that the condensates formed in LTE are less concentrated in TDP-43 than the ones created in buffer, which makes sense since the lysate contains other components that can be recruited to phase separated compartments, such as DNA, lipids, or proteins. At 2M TMAO or in LTE, the width of the peaks and therefore average diameter of the condensates increases linearly as a function of protein concentration (Figure S14). This is in agreement with the behaviour of RNA-binding proteins of the FUS-EWSR1-TAF15 family reported by Mrityunjoy et al^36^. It also aligns with the models developed by Amico et al^39^.

We also took advantage of the real-time acquisition afforded by this single molecule technique to investigate the dynamics of condensate formation and growth. Based on our data, the first nanocondensates quickly form within a minute (Figure 4). TMAO seems to have little effect on this parameter; rather, increased TMAO concentration could contribute to stabilizing these nuclei, allowing larger and brighter objects to form more rapidly. This is also partially seen during the dissolution experiments where bigger and brighter objects persist longer in the presence of TMAO (Figure 3). Within the first hour, the number of condensates in the presence of TMAO decreases while the average size (peak width) increases; the fraction of protein in the condensates (total intensity) remains unchanged (Figure S16). This can also be seen with the transition from the BL quadrant to the BR quadrant in our fingerprint analysis (Figure 5). In lysate (LTE), a similar conversion occurs more slowly (Figure S11). It is still unclear whether these changes are linked to coalescence or stabilization and growth of a subset of nanocondensates.

In conclusion, we presented a single molecule method to study the formation of nanocondensates. Our tool quantifies condensate size and concentration to obtain detailed information on the steps from initial formation of small nuclei to nanocondensates then on to micron sized droplets. As monitoring protein LLPS occurring in cells is becoming an active field of research^40–44^, we envision our method as a useful tool to support findings of nanocondensation in cells^45,46^. Our assay can be applied to different proteins and conditions ranging from buffers to crowded cellular environments. Since our technique can be expanded to detect two labels simultaneously, mechanisms of condensate formation between two proteins or proteins and RNA could be studied as well. Thus, investigating these increasingly complex systems could close the gap between observing phase separation *in vitro* and *in vivo*. Fluorescence fluctuation spectroscopy can also be applied to live cell microscopy, and it may also be possible to adapt some of the analyses presented here to study the dynamics of protein condensation in cells^14^. Our method could also be effective when studying the interplay between protein aggregation and condensation, particularly in the context of neurodegenerative disorders^47^. Increasingly, phase separation is understood as an early step in the process of protein aggregation. Our method could help distinguish between the two processes and be adapted to study the transition between liquid and solid-like states. Bridging this gap could improve the development of novel therapeutics for the regulation of protein condensation and aggregation, ultimately leading to preventing aberrant liquid-to-solid transitions^48^.

## Acknowledgments

The authors would like to thank Profs. Josephine and Allan Ferreon for the generous gift of the purified TDP-43 protein and TDP-43 construct. The cell-free lysate was produced by Dr. D J.B. Hunter (UNSW and IMB, University of Queensland); liposomes (Figure 6) were provided by Martin Do (Boecking lab, UNSW) and HVB (hepatitis virus B) capsids were provided by A.Prof D. Jacques (UNSW). The authors would also like to thank the group members for their help and support with reagents and experiments.

## Methods

### Production of Atto647N-labeled TDP-43 CTD monomers

Purified TDP-43 monomers were obtained from Profs. Allan and Josephine Ferreon, who described the first phase diagram of TDP-43 phase separation in the presence of TMAO (Choi et al, 2018)^12^. This ensures perfect agreement between previous data and our single molecule observations. Unlabelled TDP-43 CTD monomers were labelled with Atto647N-NHS dyes (ATTO-TEC GmbH) with stochiometric amount, using the manufacturer’s protocol. Purification of the labelled proteins was done by repeated filtration in a 3 kDa filter (Amicon Ultra-15 Centrifugal Filter Unit, Merck).

### Single molecule experiments

Samples obtained from expression and purification, as described above, were loaded into a custom-made silicone well plate with a 70 x 80 mm glass coverslip (ProSciTech, Kirwan, QLD, Australia). In most experiments αβγ buffer (10 mM sodium acetate, 10 mM sodium phosphate, 10 mM glycine, 200 mM NaCl, pH 7.5) was used in the presence of different concentrations of trimethylamine *N*-oxide (TMAO; 0, 0.5, 1, 1.5, and 2M)^12^. Samples are prepared individually by adding 2 µL of Atto647N-labeled TDP-43 LCD (5 µM) to 16 µL of αβγ buffer and TMAO At a given concentration. After addition of the labeled protein, the measurements were started. After ∼50 seconds, the optical path going towards the detector was blocked (by switching the knob between the eye piece and the detector path) and 2 µL unlabeled TDP-43 LCD was added to the sample and mixed thoroughly to ensure homogeneity. The measurement resumed after addition of the unlabeled protein. Plates were analysed at room temperature on Zeiss Axio Observer microscope (Zeiss, Oberkochen, Germany) with a custom-built data acquisition set-up. Illumination is provided by a 488 nm laser beam focussed in the sample volume using a ×40 magnification, 1.2 Numerical Aperture water immersion objective (Zeiss, Oberkochen, Germany). This creates a very small observation volume in solution (∼1 fL), through which fluorescent proteins diffuse. Light emitted by the fluorophores is transmitted through a 560 nm dichroic mirror and a 525/50 nm band-pass filter before being detected by a photon counting detector (Micro Photon Devices, Bolzano, Italy). Photons are recorded in 1ms time bins and analysed using Lab-VIEW 2018 version 18.0 (National Instruments)^49^. Each sample is measured for 10 minutes and traces were analyzed using a custom python script on Spyder version 5.4.3 (see Supplementary Information).

The “single molecule analysis” script reports on the following, for each peak: 1. The prominence of each peak which corresponds to the intensity of a peak corrected for background fluorescence. 2. The residence time or the peak width of a burst. Conservatively, it is defined as the difference of two time points in which the intensity values are at 50% of the prominence. 3. The total intensity of each fluorescence burst, defined as the sum of intensities during the peak width to estimate the area under the curve (AUC). We observed that calculating peak width at 50% of prominence provided a more reliable metric compared to using 90% of prominence or 100%, even if this slightly underestimates peak duration and total area under the curve.

For each trace, the software generates a CSV file containing these 3 values for each peak. In addition, the script generates a summary file (final_analyse.csv), with the list of the average values for residence times, peak intensity, AUC, and the sum of the AUCs for all peaks for all the traces.

### Quadrant analysis python script to fingerprint the condensates

The “quadrant analysis” python script uses the output CSV files from the peak analysis. The ‘fingerprints’ of the condensates can be quantified by plotting the peak intensity and peak width obtained for each peak. To define 4 sub-quadrants as shown in Figure 5A, thresholds for the peak width and intensity must be determined (see Supplementary Information). The script generates a CSV file with the percentage and absolute number of peaks in each quadrant for each file.

### Hexanediol experiments

Stock solutions of 1,6-hexanediol (Sigma-Aldrich ref 240117) were prepared in ddH2O with concentrations in w/v% (10, 20, 30, 40, 50). Again, samples are prepared individually by adding 12 µL of 1x αβγ buffer with 2M TMAO, 2 µL of 5 µM Atto647N-labeled TDP-43 LCD, and 2 µL of 5 µM unlabeled TDP-43 LCD. To ensure LLPS is occurring in each sample a positive control measurement of 5 minutes is done before adding 1,6-hexanediol. If phase separation can be observed the measurement is continued for 20 minutes after addition of 4 µL 1,6-hexanediol at the desired concentration (final concentrations are 0, 2, 4, 6, 8, and 10 w/v% 1,6-hexanediol, 500 nM Atto647N-labeled TDP-43 LCD, and 500 nM unlabeled TDP-43 LCD).

### Dilution experiments

To determine reversibility of LLPS we used a slightly different sample preparation method compared to the general single molecule experiments. Initially, a sample is prepared identically to the general method by to 16 µL of 1x αβγ buffer ((10 mM sodium acetate, 10 mM sodium phosphate, 10 mM glycine, 200 mM NaCl, pH 7.5) with 2M TMAO, 2 µL of 5 µM Atto647N-labeled TDP-43 LCD, and 2 µL of 15 µM unlabeled TDP-43 LCD. Subsequent samples are prepared by diluting the previous sample in a dilution buffer (1x αβγ buffer with 2M TMAO containing 500 nM Atto647N-labeled TDP-43 LCD) to reach final concentrations of 2000, 1750, 1500, 1250, 1000, and 750 nM TDP-43 LCD. In these samples the concentration of the labeled protein is kept constant at 500 nM. Each sample is measured for 10 minutes.

### Kymograph analysis

The frame analysis was developed to split measurements into specific time frames of interest. This functionality was integrated into the initial custom python analysis script which automatically determines the results for each time frame instead of the full trace. The script splits the data into time frames. The number of frames can be changed by altering the value for the variable “number_of_frames” in the script. As an example, if a fluorescent trace was measured for 600000 ms and “number_of_frames” is set to 10. The fluorescent trace will be split into 10 frames of 60000 ms each. After running the script, another CSV file will be created which contains the peak number, mean peak intensity, mean peak width, mean peak integral, and total intensity for each frame for every fluorescence trace given.

### First peak analysis

The time to the first condensate formation is represented by the peak position of the first peak and can be found in the data files after running the peak analysis script. Furthermore, a new custom python script, “first peak analysis” was created to automatically determine the peak position (ms) of the first peak above a certain intensity for each file. In case there is no peak above a certain intensity, a value of 600000 is given (for a 10-minute trace). A new CSV file will be created which lists the peak positions of the first peak for a range of intensity values between 0 and 20000 with increments of 500.

### Size calibration experiments

Commercial fluorescent beads of 430, 200 and 100 nm diameter were used as well as liposomes (120 nm) and purified Hepatitis B capsids (45 nm) to generate a calibration curve (see Supplementary Table). Beads were diluted in αβγ buffer (except for liposomes) to a final concentration that allowed the detection of 50-200 events per trace. Several 5 minutes fluorescence traces were acquired as described above. Data was analyzed using the custom python peak analysis. A calibration curve was obtained by plotting the mean peak width against the physical diameter of the beads and fitted by a linear regression.

## Abrreviations

TDP-43: TAR DNA-binding protein 43
TAR: Trans-activation response element
LLPS: Liquid-liquid phase separation
LCD: low complexity domain
TMAO: Trimethylamine N-oxide
LTE: Leishmania tarentolae extract
TL: Top left
TR: Top right
BL: Bottom left
BR: Bottom right
FUS: Fused in sarcoma
cGAS: cyclic GMP–AMP synthase

## Bibliography

1. Wang, B. et al. Liquid–liquid phase separation in human health and diseases. Signal Transduction and Targeted Therapy vol. 6 Preprint at 10.1038/s41392-021-00678-1 (2021).

2. Gomes, E. & Shorter, J. The molecular language of membraneless organelles. Journal of Biological Chemistry vol. 294 7115–7127 Preprint at 10.1074/jbc.TM118.001192 (2019).

3. Brangwynne, C. P., Tompa, P. & Pappu, R. V. Polymer physics of intracellular phase transitions. Nat Phys 11, 899–904 (2015).

4. Jin, X., et al. Membraneless Organelles Formed by Liquid-Liquid Phase Separation Increase Bacterial Fitness. Sci. Adv vol. 7 (2021).

5. Alberti, S., Gladfelter, A. & Mittag, T. Considerations and Challenges in Studying Liquid-Liquid Phase Separation and Biomolecular Condensates. Cell vol. 176 419–434 Preprint at 10.1016/j.cell.2018.12.035 (2019).

6. Banani, S. F., Lee, H. O., Hyman, A. A. & Rosen, M. K. Biomolecular condensates: Organizers of cellular biochemistry. Nature Reviews Molecular Cell Biology vol. 18 285–298 Preprint at 10.1038/nrm.2017.7 (2017).

7. Mondal, S. et al. Multivalent interactions between molecular components involved in fast endophilin mediated endocytosis drive protein phase separation. Nat Commun 13, (2022).

8. Dignon, G. L., Best, R. B. & Mittal, J. Annual Review of Physical Chemistry Biomolecular Phase Separation: From Molecular Driving Forces to Macroscopic Properties. Annu Rev Phys Chem 71, 53–75 (2020).

9. Toledo, P. L., Gianotti, A. R., Vazquez, D. S. & Ermácora, M. R. Protein nanocondensates: the next frontier. Biophysical Reviews vol. 15 515–530 Preprint at 10.1007/s12551-023-01105-1 (2023).

10. Keber, F. C., Nguyen, T., Mariossi, A., Brangwynne, C. P. & Wühr, M. Evidence for widespread cytoplasmic structuring into mesoscale condensates. Nat Cell Biol 26, 346–352 (2024).

11. Gambin, Y. et al. Single-Molecule Fluorescence Reveals the Oligomerization and Folding Steps Driving the Prion-like Behavior of ASC. J Mol Biol 430, 491–508 (2018).

12. Choi, K. J. et al. A Chemical Chaperone Decouples TDP-43 Disordered Domain Phase Separation from Fibrillation. Biochemistry 57, 6822–6826 (2018).

13. Joachim D. Mueller. Fluorescence Fluctuation Spectroscopy. Encyclopedia of Biophysics 800–803 (2013) doi:10.1007/978-3-642-16712-6.

14. Priest, D. G., Solano, A., Lou, J. & Hinde, E. Fluorescence fluctuation spectroscopy: An invaluable microscopy tool for uncovering the biophysical rules for navigating the nuclear landscape. Biochemical Society Transactions vol. 47 1117–1129 Preprint at 10.1042/BST20180604 (2019).

15. Brown, J. W. P. et al. Single-molecule detection on a portable 3D-printed microscope. Nat Commun 10, (2019).

16. Lau, D. et al. Single Molecule Fingerprinting Reveals Different Amplification Properties of α-Synuclein Oligomers and Preformed Fibrils in Seeding Assay. ACS Chem Neurosci 13, 883–896 (2022).

17. O’Carroll, A. et al. Pathological mutations differentially affect the self-assembly and polymerisation of the innate immune system signalling adaptor molecule MyD88. BMC Biol 16, (2018).

18. Sierecki, E. et al. Nanomolar oligomerization and selective co-aggregation of α-synuclein pathogenic mutants revealed by single-molecule fluorescence. Sci Rep 6, (2016).

19. Jo, M. et al. The role of TDP-43 propagation in neurodegenerative diseases: integrating insights from clinical and experimental studies. Experimental and Molecular Medicine vol. 52 1652–1662 Preprint at 10.1038/s12276-020-00513-7 (2020).

20. Huang, W. et al. TDP-43: From Alzheimer’s Disease to Limbic-Predominant Age-Related TDP-43 Encephalopathy. Frontiers in Molecular Neuroscience vol. 13 Preprint at 10.3389/fnmol.2020.00026 (2020).

21. Nelson, P. T. et al. Limbic-predominant age-related TDP-43 encephalopathy (LATE): Consensus working group report. Brain vol. 142 1503–1527 Preprint at 10.1093/brain/awz099 (2019).

22. Steinacker, P., Barschke, P. & Otto, M. Biomarkers for diseases with TDP-43 pathology. Molecular and Cellular Neuroscience vol. 97 43–59 Preprint at 10.1016/j.mcn.2018.10.003 (2019).

23. Babinchak, W. M. et al. The role of liquid-liquid phase separation in aggregation of the TDP-43 low-complexity domain. Journal of Biological Chemistry 294, 6306–6317 (2019).

24. François-Moutal, L. et al. Structural Insights Into TDP-43 and Effects of Post-translational Modifications. Frontiers in Molecular Neuroscience vol. 12 Preprint at 10.3389/fnmol.2019.00301 (2019).

25. Chris, A., Ferreon, M., Moosa, M., Gambin, Y. & Deniz, A. A. Counteracting chemical chaperone effects on the single-molecule α-synuclein structural landscape. PNAS 109, 17826–17831 (2012).

26. Hunter, D. J. B., Bhumkar, A., Giles, N., Sierecki, E. & Gambin, Y. Unexpected instabilities explain batch-to-batch variability in cell-free protein expression systems. Biotechnol Bioeng 115, 1904–1914 (2018).

27. Bandyopadhyay, A. et al. Chemical chaperones assist intracellular folding to buffer mutational variations. Nat Chem Biol 8, 238–245 (2012).

28. Garg, D. K. & Bhat, R. Modulation of assembly of TDP-43 low-complexity domain by heparin: From droplets to amyloid fibrils. Biophys J 121, 2568–2582 (2022).

29. Polyansky, A. A., Gallego, L. D., Efremov, R. G., Köhler, A. & Zagrovic, B. Protein compactness and interaction valency define the architecture of a biomolecular condensate across scales. Elife 12, (2023).

30. Sanchez-Burgos, I., Joseph, J. A., Collepardo-Guevara, R. & Espinosa, J. R. Size conservation emerges spontaneously in biomolecular condensates formed by scaffolds and surfactant clients. Sci Rep 11, (2021).

31. Yokosawa, K. et al. Quantification of the concentration in a droplet formed by liquid–liquid phase separation of G-quadruplex-forming RNA. Chem Phys Lett 826, (2023).

32. Keating, S. S., Bademosi, A. T., San Gil, R. & Walker, A. K. Aggregation-prone TDP-43 sequesters and drives pathological transitions of free nuclear TDP-43. Cellular and Molecular Life Sciences 80, (2023).

33. Haider, R., Penumutchu, S., Boyko, S. & Surewicz, W. K. Phosphomimetic substitutions in TDP-43’s transiently α-helical region suppress phase separation. Biophys J (2024) doi:10.1016/j.bpj.2024.01.001.

34. Vazquez, D. S., Toledo, P. L., Gianotti, A. R. & Ermácora, M. R. Protein conformation and biomolecular condensates. Current Research in Structural Biology vol. 4 285–307 Preprint at 10.1016/j.crstbi.2022.09.004 (2022).

35. Babinchak, W. M. et al. Small molecules as potent biphasic modulators of protein liquid-liquid phase separation. Nat Commun 11, (2020).

36. Ray, S. et al. Mass photometric detection and quantification of nanoscale α-synuclein phase separation. Nat Chem 15, 1306–1316 (2023).

37. Kar, M., et al. Phase-separating RNA-binding proteins form heterogeneous distributions of clusters in subsaturated solutions. PNAS 119, (2022).

38. Yao, Y., Wang, W. & Chen, C. Mechanisms of phase-separation-mediated cGAS activation revealed by dcFCCS. PNAS Nexus 1, (2022).

39. Amico, T. et al. Physics of Living Systems A scale-invariant log-normal droplet size distribution below the transition concentration for protein phase separation. Elife 13, (2024).

40. Jun-Yan Kang et al. LLPS of FXR1 drives spermiogenesis by activating translation of stored mRNAs. Science (1979) 337, (2022).

41. Wegmann, S. et al. Tau protein liquid–liquid phase separation can initiate tau aggregation. EMBO J 37, (2018).

42. Maurel, C. et al. The post-translational modification SUMO affects TDP-43 phase separation, compartmentalization, and aggregation in a zebrafish model. doi:10.1101/2022.08.14.503569.

43. Zhu, S. et al. Liquid-liquid phase separation of RBGD2/4 is required for heat stress resistance in Arabidopsis. Dev Cell 57, 583–597.e6 (2022).

44. Forman-Kay, J. D., Ditlev, J. A., Nosella, M. L. & Lee, H. O. What are the distinguishing features and size requirements of biomolecular condensates and their implications for RNA-containing condensates? RNA 28, 36–47 (2022).

45. Owyong, T. C., et al. A General Fluorescence-Based Method for Quantifying and Mapping Biomolecular Polarity In Vitro and In Cells. BioRxiv (2023) doi:10.1101/2023.02.07.526546.

46. Zhang, S., Hinde, E., Parkyn Schneider, M., Jans, D. A. & Bogoyevitch, M. A. Nuclear bodies formed by polyQ-ataxin-1 protein are liquid RNA/protein droplets with tunable dynamics. Sci Rep 10, (2020).

47. Chakraborty, P. & Zweckstetter, M. Role of aberrant phase separation in pathological protein aggregation. Current Opinion in Structural Biology vol. 82 Preprint at 10.1016/j.sbi.2023.102678 (2023).

48. Mitrea, D. M., Mittasch, M., Gomes, B. F., Klein, I. A. & Murcko, M. A. Modulating biomolecular condensates: a novel approach to drug discovery. Nat Rev Drug Discov 21, 841–862 (2022).

49. Leitão, A. D. G., et al. Selectivity of Lewy body protein interactions along the aggregation pathway of α-synuclein. Commun Biol 4, (2021).

