## Supplementary information for "Quantification of nanocondensates formation at the single molecule level"

**Supplementary information: Quantification of nanocondensates formation at the single molecule level**

**Figure S1. Domain composition and structure of the human TDP-43**
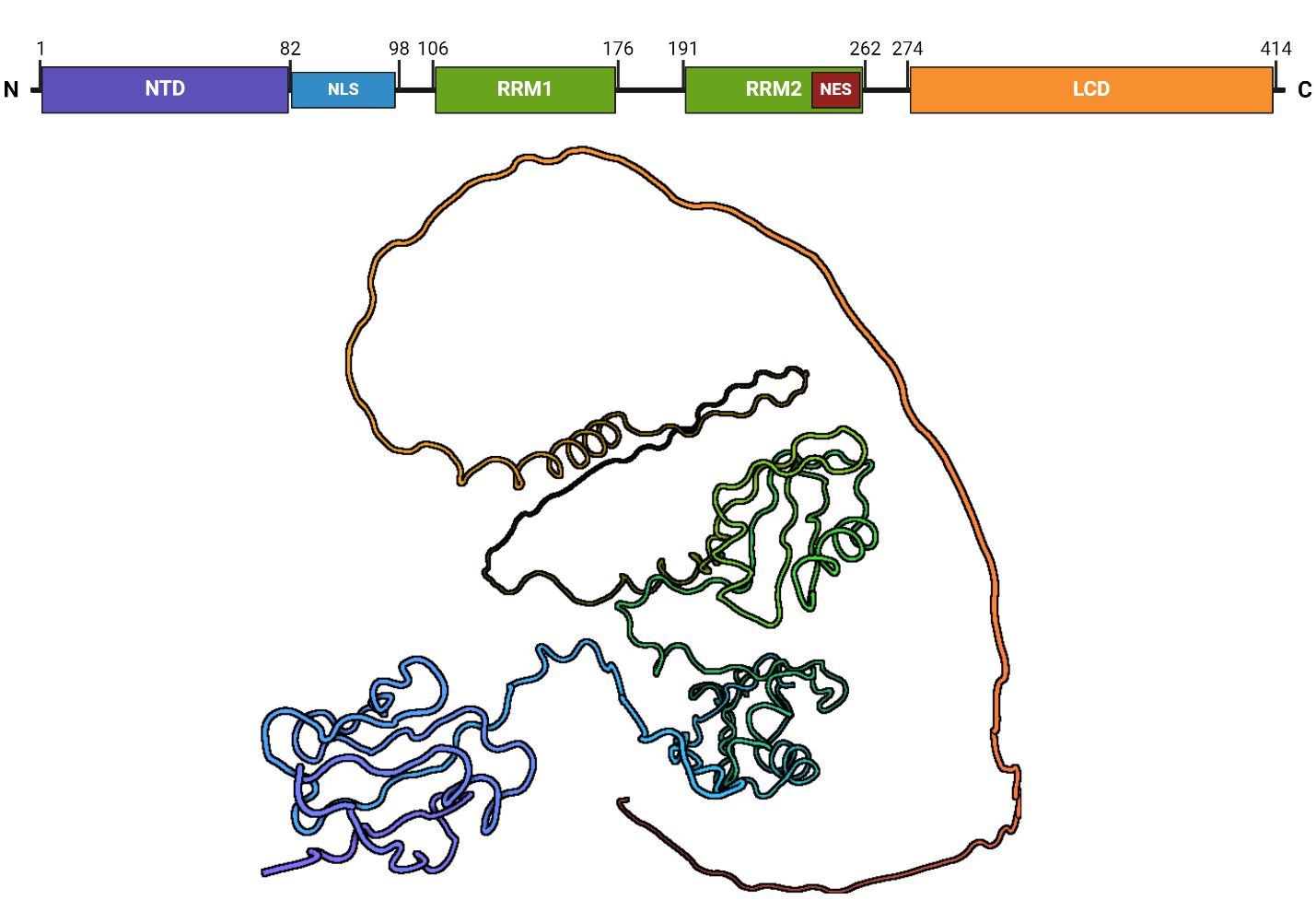

Schematic overview of the human TDP-43 proteins with the domains and corresponding amino acid numbers indicated. The depicted domains include the N-terminal domain (NTD in purple), the nuclear localization signal (NLS in blue), the two RNA-recognition motifs (RRM1 and RRM2 in green) including a nuclear export signal (NES in red) within RRM2, and the low-complexity domain (LCD in orange). Structure of human TDP-43 predicted by Alphafold (PDB: Q13148). Colors correspond to the domains shown above.

**Figure S2. Individual replicates of TMAO titration experiments**

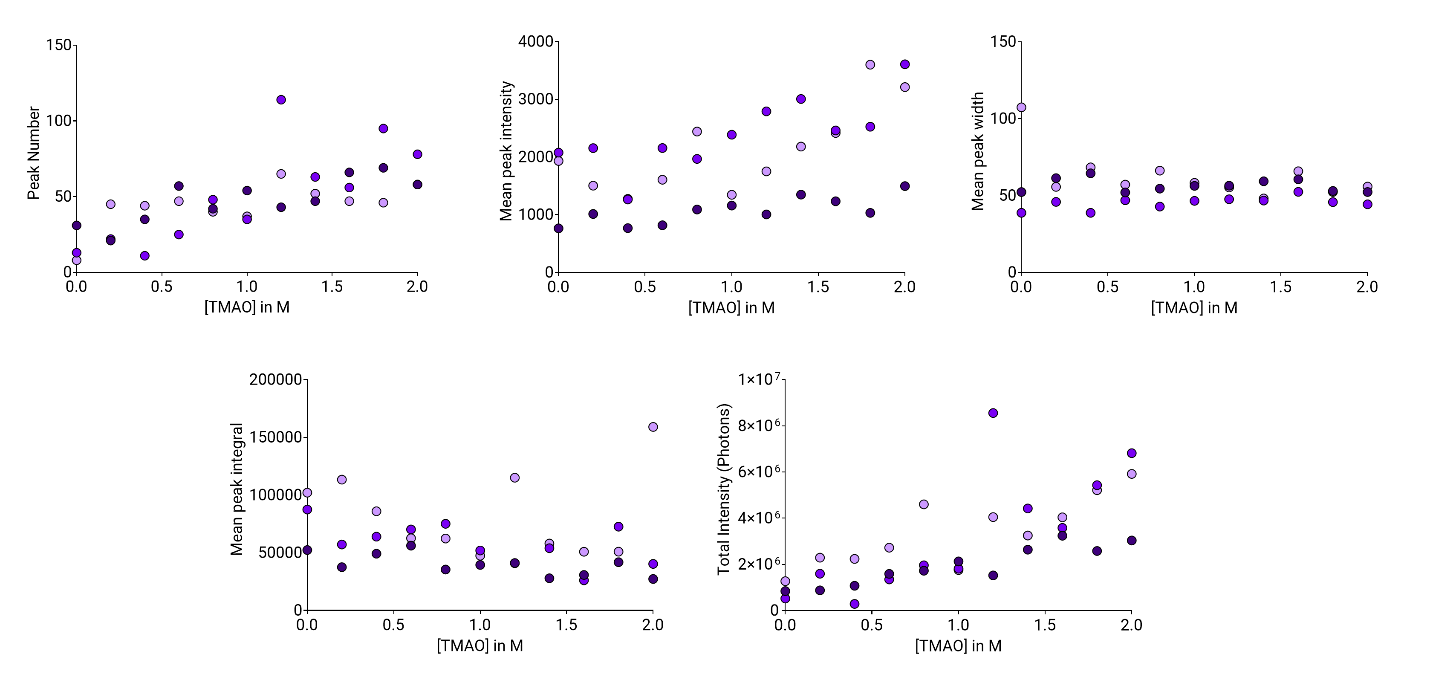

Graphs showing the peak number, mean peak intensity, mean peak width, mean peak integral, and total intensity of individual replicates for TMAO titration experiments conducted in αβγ buffer. Each replicate is represented by a different shade of purple.

**Figure S3.** **Effect of** **TMAO concentration**

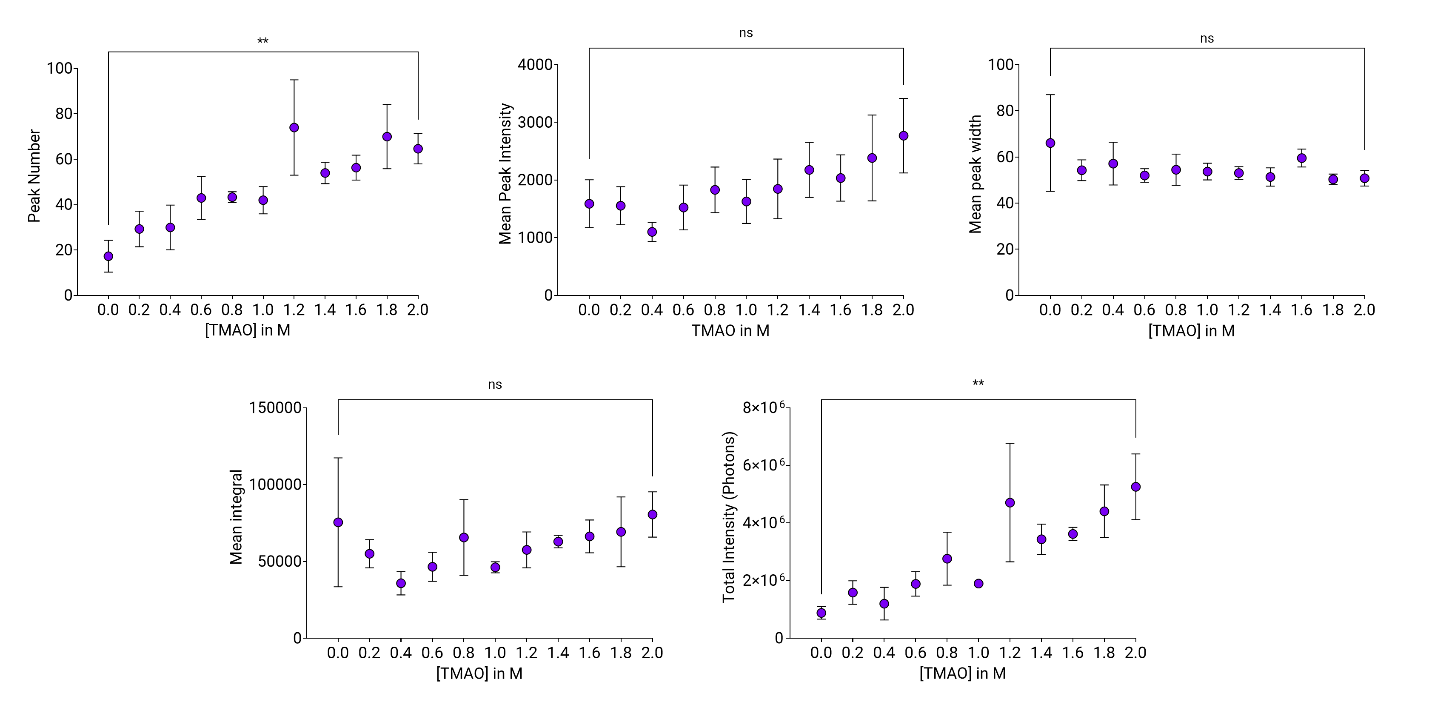

Graphs depicting the peak number, mean peak intensity, mean peak width, mean peak integral, and total intensity as a function of TMAO concentration, obtained from three separate titration experiments (see Figure S2). In all graphs, the mean is presented, and the error bars are standard error of the mean (SEM). Statistical analysis was conducted using the Friedman test, with significance levels denoted as follows: p ≤ 0.05 (*), p ≤ 0.01 (**), p ≤ 0.001 (***), p ≤ 0.0001 (****), and nonsignificant (n.s.).

**Figure S4. Fluorescence time traces of condensate formation in 2M TMAO and LTE**
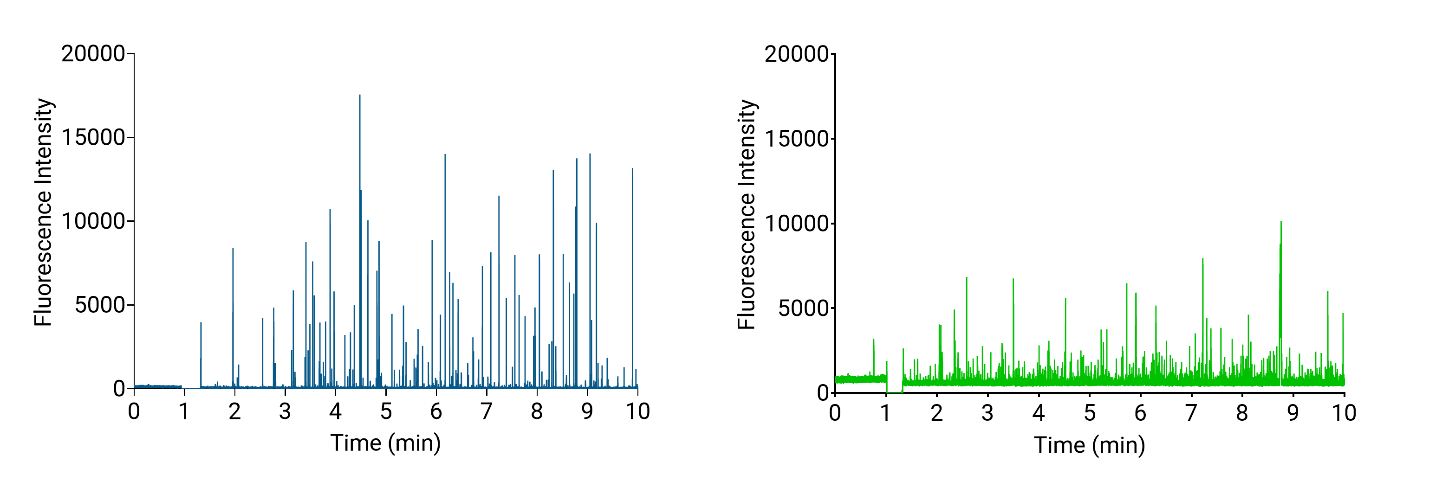
Representative fluorescence intensity traces (in photon per ms) obtained for 500 nM Atto647N-labeled TDP-43 LCD and 500 nM unlabeled TDP-43 LCD in 2M TMAO (blue) or in LTE (green).

**Figure S5. TDP-43 LCD titration experiments in 2M TMAO or LTE**

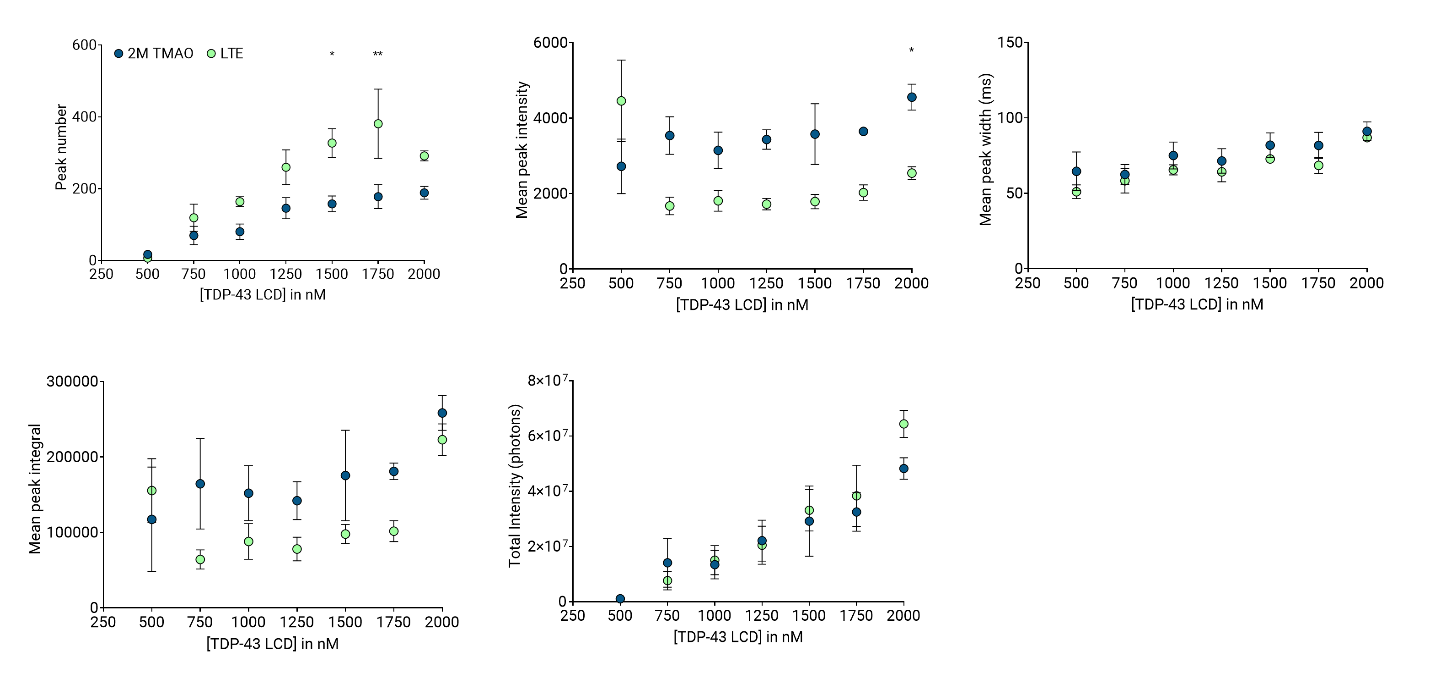
Graphs presenting the peak number, mean peak intensity, mean peak width, mean peak integral, and total intensity as a function of TDP-43 LCD concentration, obtained from three separate titration experiments, either in 2M TMAO (blue) or LTE (green). In all graphs, the mean was calculated, and the error bars are standard error of the mean (SEM) and statistics used Friedman test, p ≤ 0.05 (*), p ≤ 0.01 (**), p ≤ 0.001 (***), p ≤ 0.0001 (****), nonsignificant (n.s.).

**Figure S6. Intensity and width profiles of individual peaks in 2M TMAO and LTE**

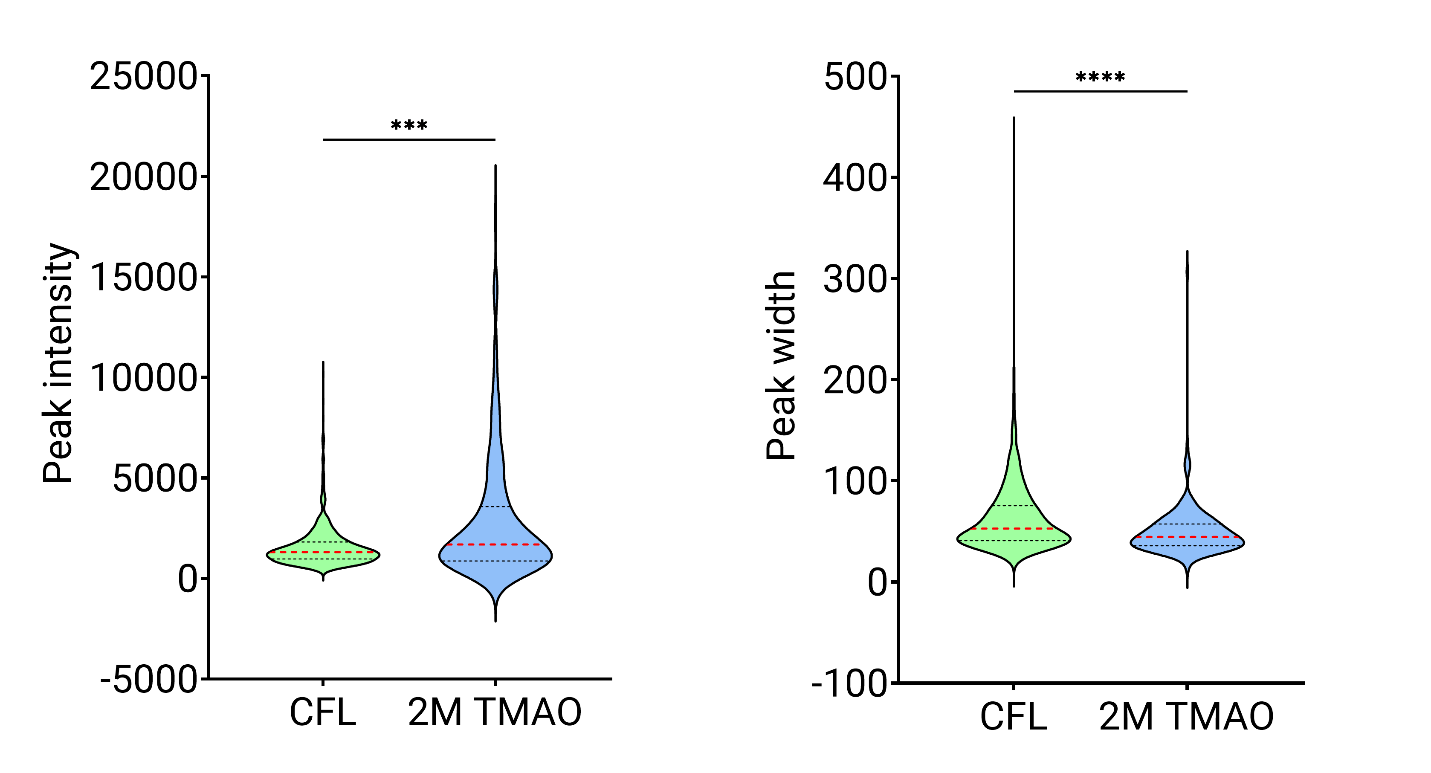

Violin plots showing the peak intensity and peak width of individual peaks from three individual measurements conducted either in 2M TMAO (blue) or cell-free lysate (LTE) (green). The red dotted lines represent the medians, and the black dotted lines indicate quartiles. Statistics used Mann-Whitney test, p ≤ 0.05 (*), p ≤ 0.01 (**), p ≤ 0.001 (***), p ≤ 0.0001 (****), nonsignificant (n.s.).

**Figure S7. Intensity of individual peaks at different 1,6-hexanediol concentrations**

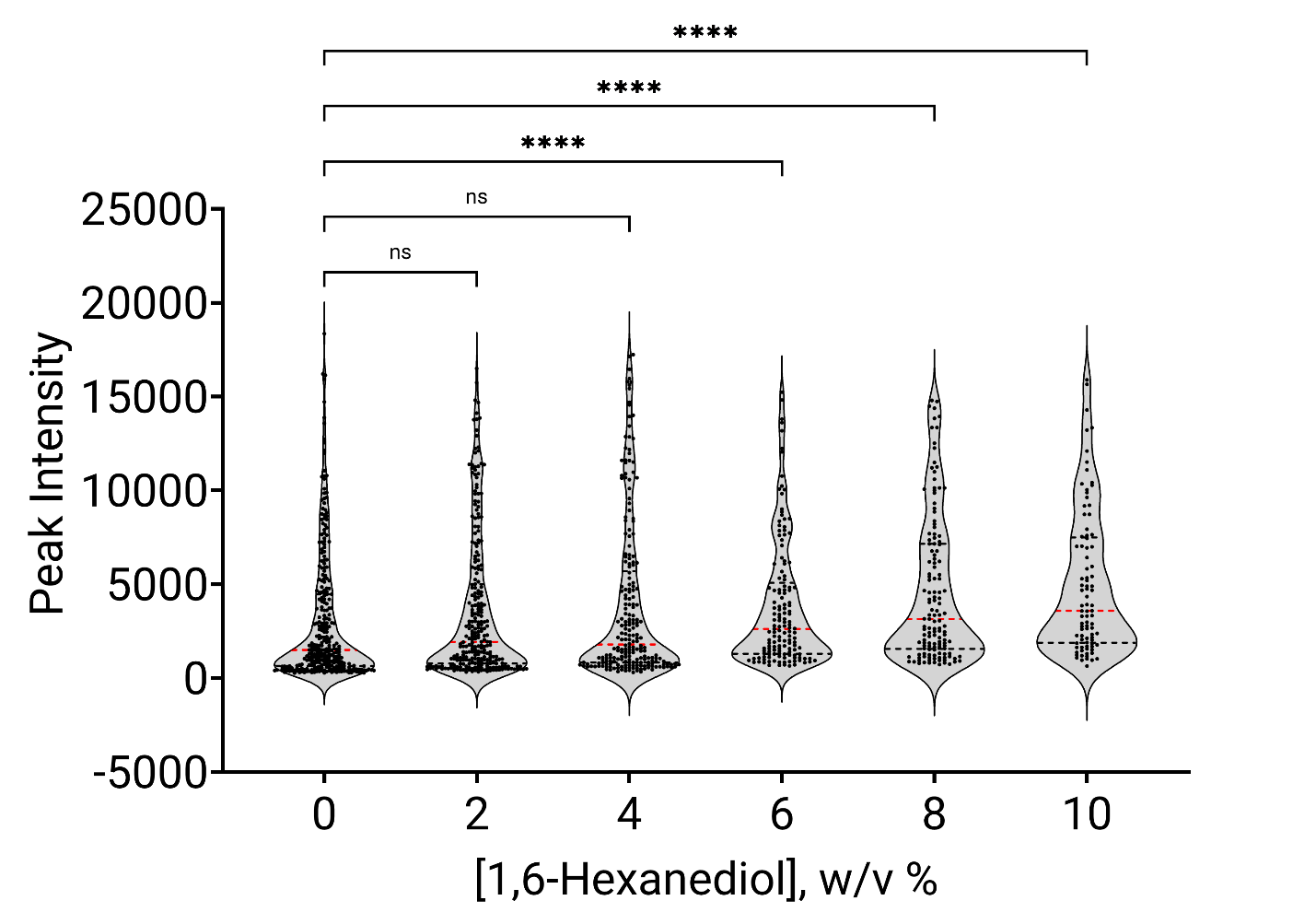

Violin plot showing the peak intensity of individual peaks (black dots) of three individual 10-minute measurements in 2M TMAO at different concentrations of 1,6-hexanediol. The red dotted lines represent the medians, and the black dotted lines indicate quartiles. Statistics used Kruskal-Wallis test, p ≤ 0.05 (*), p ≤ 0.01 (**), p ≤ 0.001 (***), p ≤ 0.0001 (****), nonsignificant (n.s.).

**Figure S8. Comparison of titration versus serial dilution in LTE**

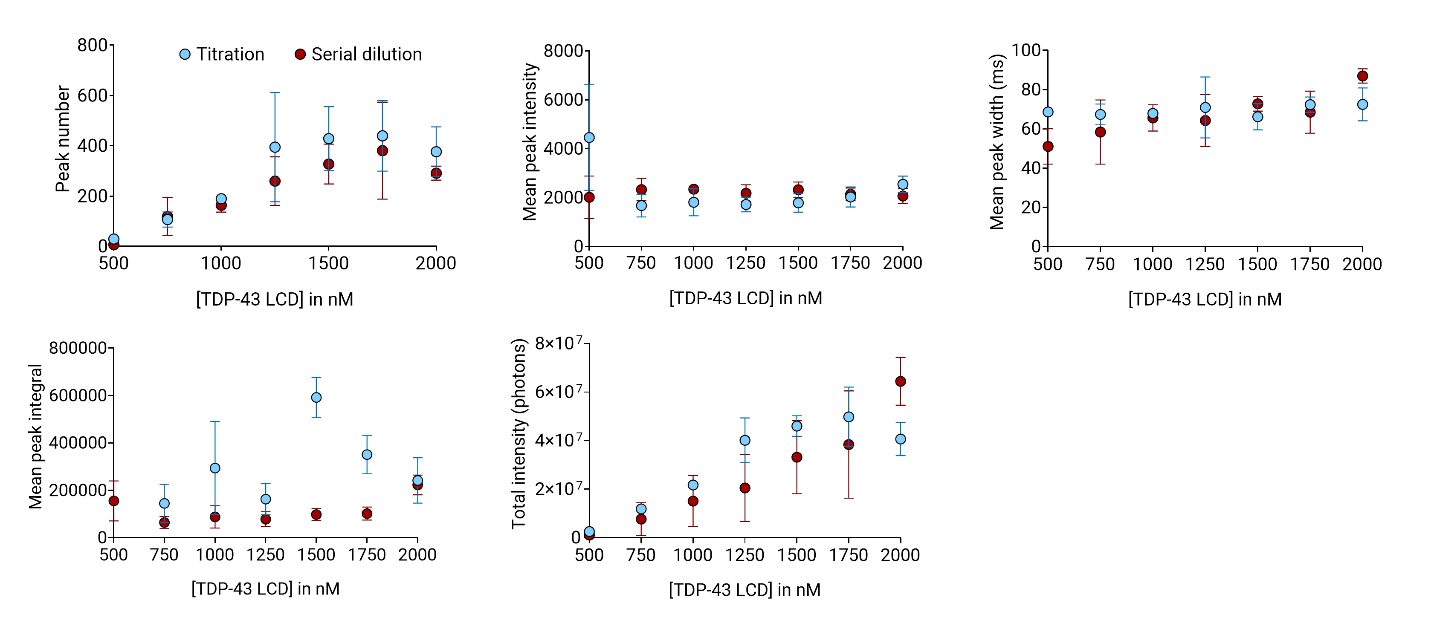
Graphs showing the peak number, mean peak intensity, mean peak width, mean peak integral, and total intensity of three measurements in LTE. The titration is indicated in blue, and the serial dilution is indicated in red. In all graphs the error bars are standard error of the mean (SEM) and statistics used two-way ANOVA (Sidak’s multiple comparisons test), all differences were nonsignificant.

**Figure S9. First peak analysis determination**
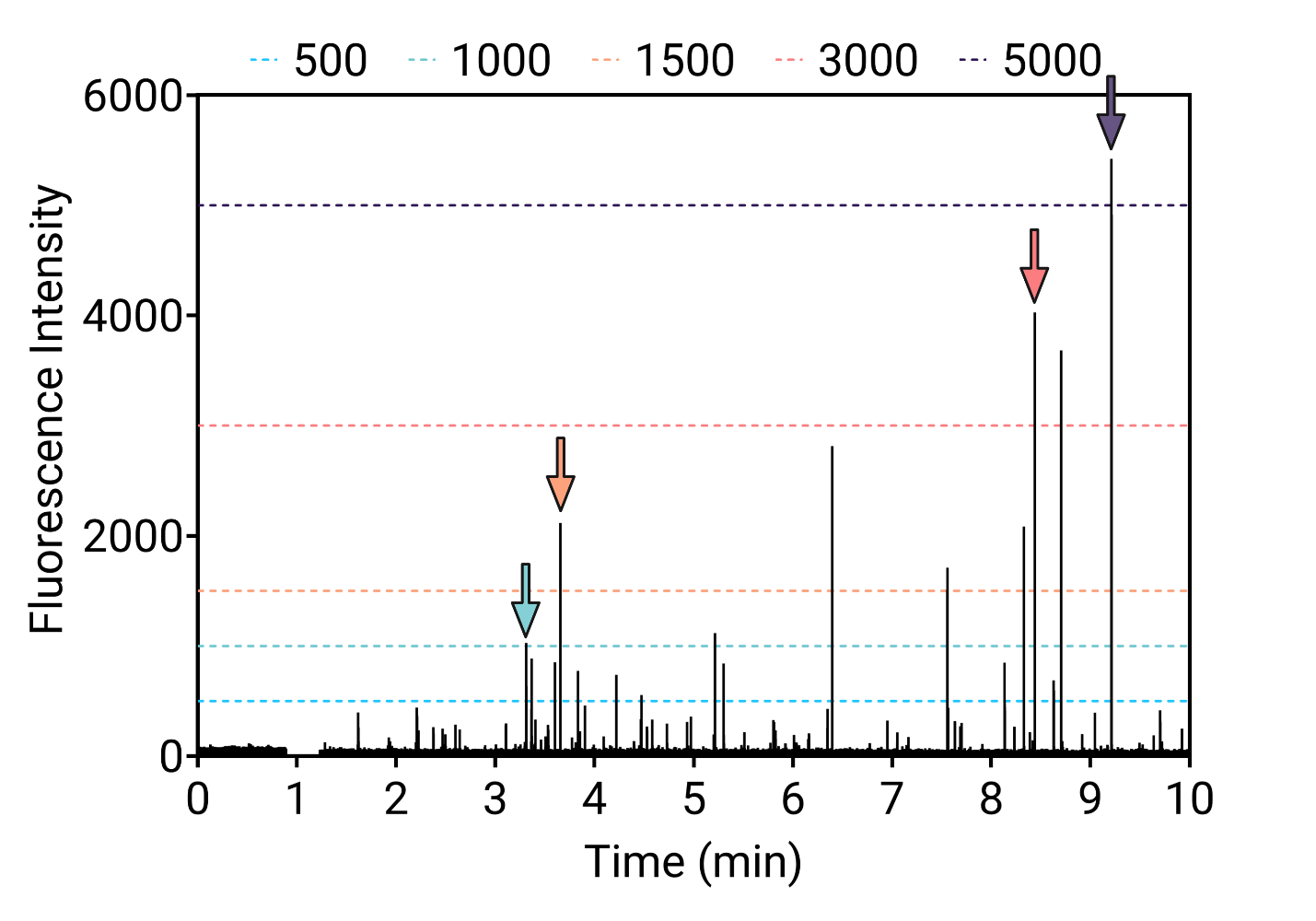

Explanatory graph displaying a fluorescence intensity trace with various horizontal dashed lines for a range of intensity thresholds. The first peak above a specific intensity is indicated by an arrow in a color corresponding to the threshold: blue (500 photons/ms), green (1000 photons/ms), orange (1500 photons/ms), red (3000 photons/ms), and purple (5000 photons/ms).

**Figure S10. Fingerprint changes upon increasing the concentration of protein**

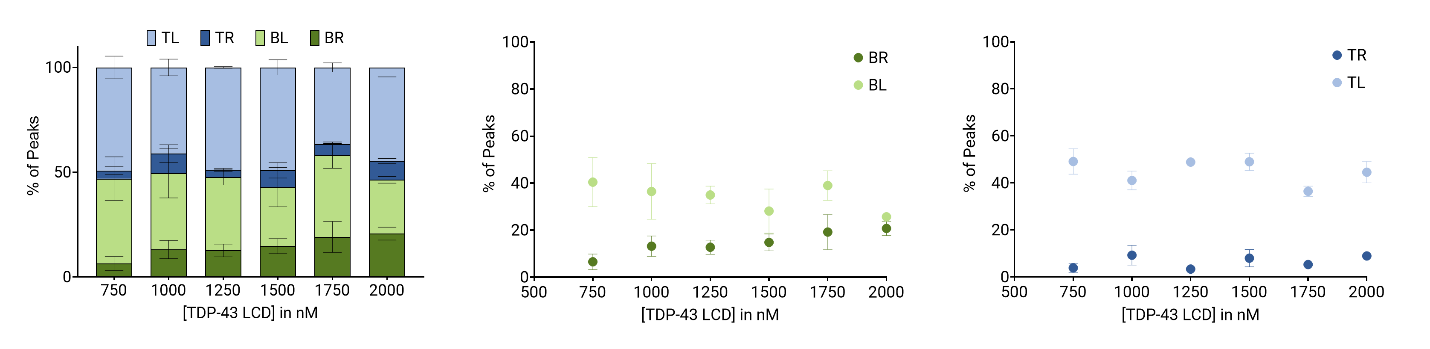

(left) Bar graph showing the percentage of peaks in each quadrant upon varying TDP-43 LCD concentration in 2M TMAO. The bottom left (BL), bottom right (BR), top left (TL), and top right (TR) quadrants are represented in light green, dark green, light blue, and dark blue respectively. (middle) Comparison between the bottom quadrants. (right) Comparison between the top quadrants. In all graphs the mean is presented, and the error bars are standard error of the mean (SEM); statistics used Kruskal-Wallis test, p ≤ 0.05 (*), p ≤ 0.01 (**), p ≤ 0.001 (***), p ≤ 0.0001 (****), nonsignificant (n.s.).

**Figure S11. Fingerprint changes over time in 2M TMAO and LTE**

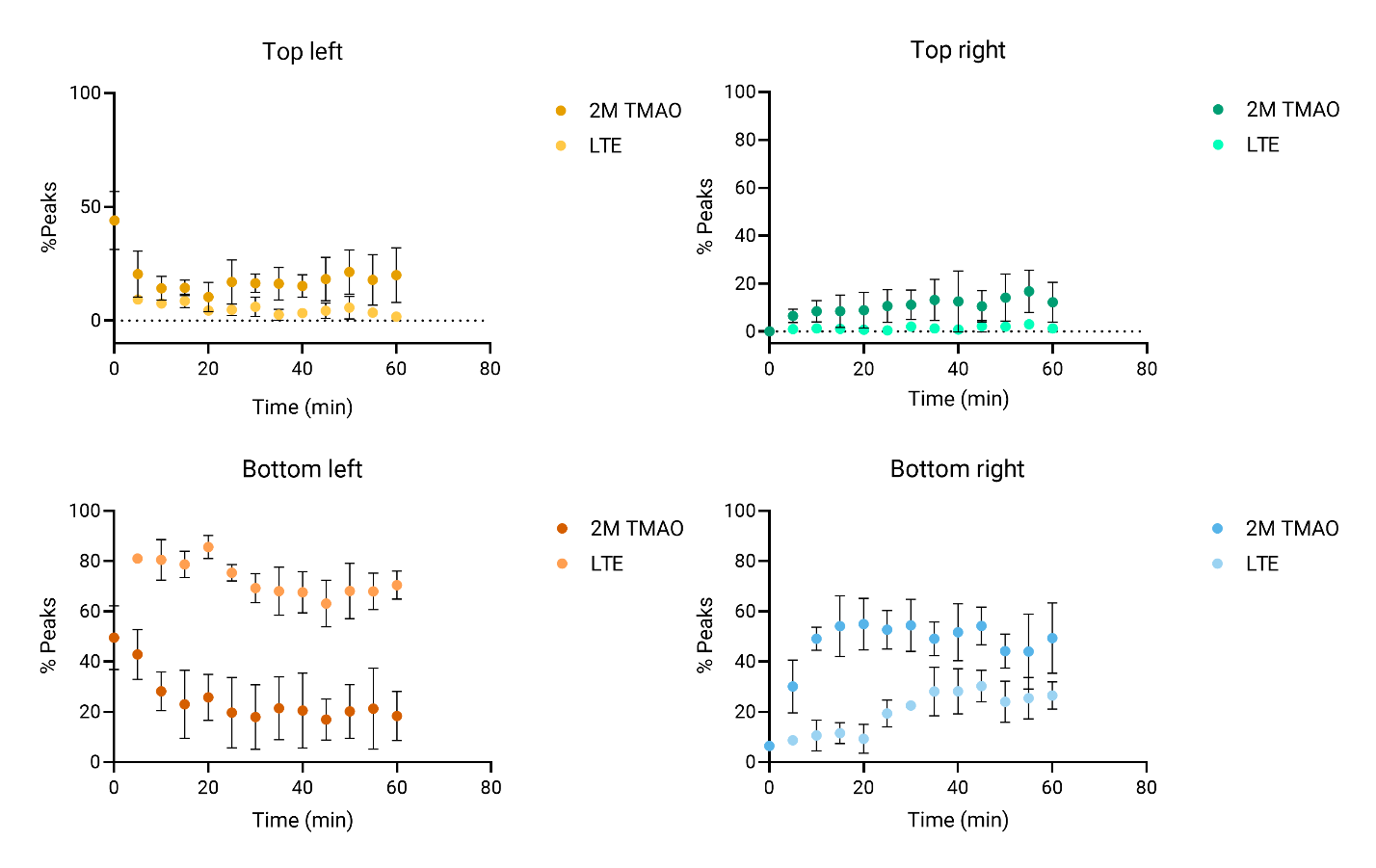

Graphs showing the percentage of peaks in each quadrant asa function of time, for 1-hour measurements in 2M TMAO and LTE (dark and light color respectively). The quadrant analysis for data acquired in samples with 2M TMAO was performed using a fixed threshold (100 peak width, and 2000 peak intensity) whereas the data acquired in LTE was analyzed with 90% thresholds using the data measured in the first 5 minutes as reference (see Methods). In all graphs the mean is presented, and error bars are standard error of the mean (SEM).

**Figure S12. Effect of normalization on calibration curves**
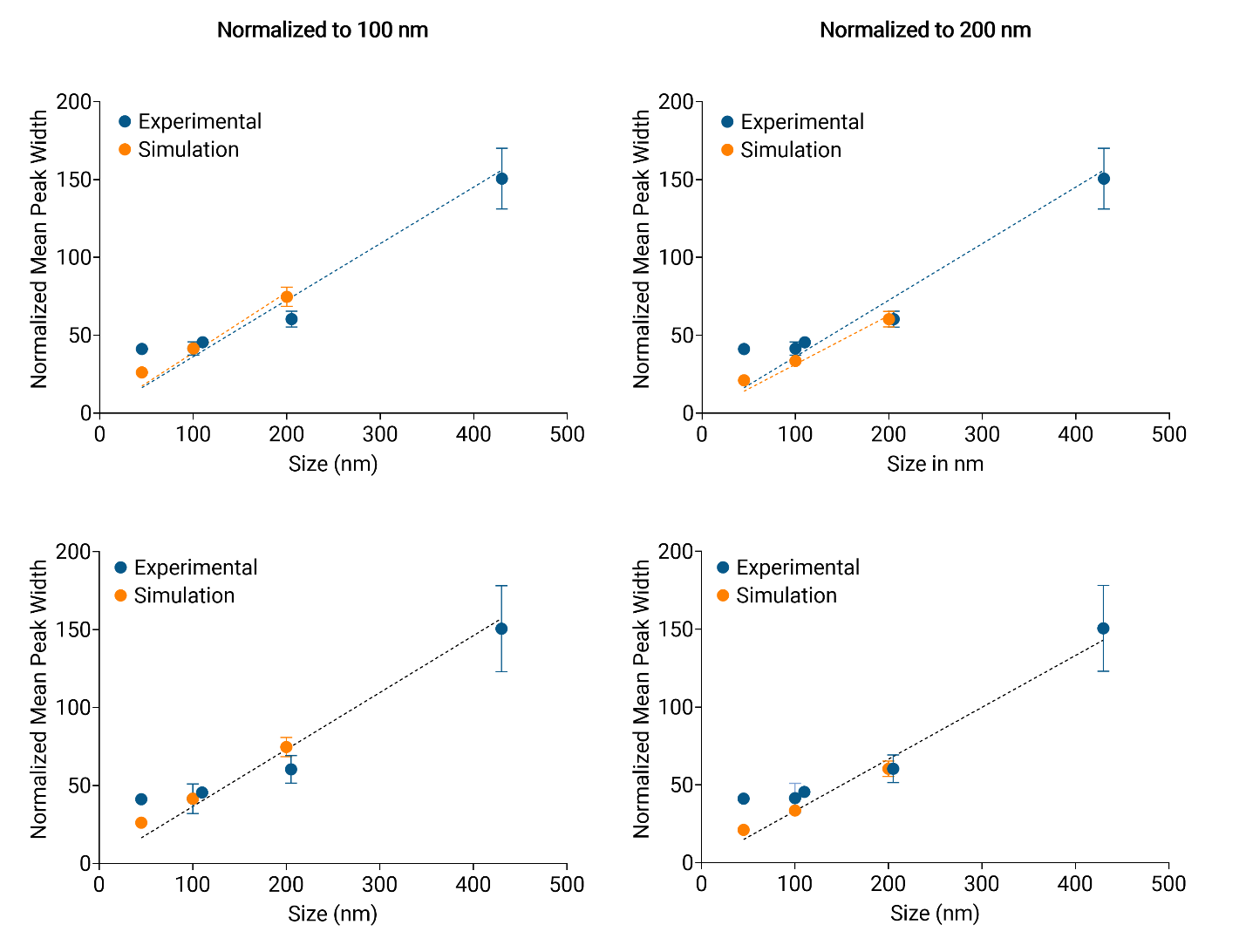

Calibration curves determined by either experimental data (blue) or simulated data (orange) using two normalizations. The graphs on the left show the simulated data normalized to the experimental data of beads with a diameter of 100 nm while the graphs on the right show the results normalized to the experimental data of beads with a diameter of 200 nm The top graphs show the fitted linear curves for the experimental and simulated data separately. The bottom graphs depict the linear fit for all data points excluding the experimental results for a size of 45 nm.

**Figure S13. Combined effect of TDP-43 and TMAO concentrations**

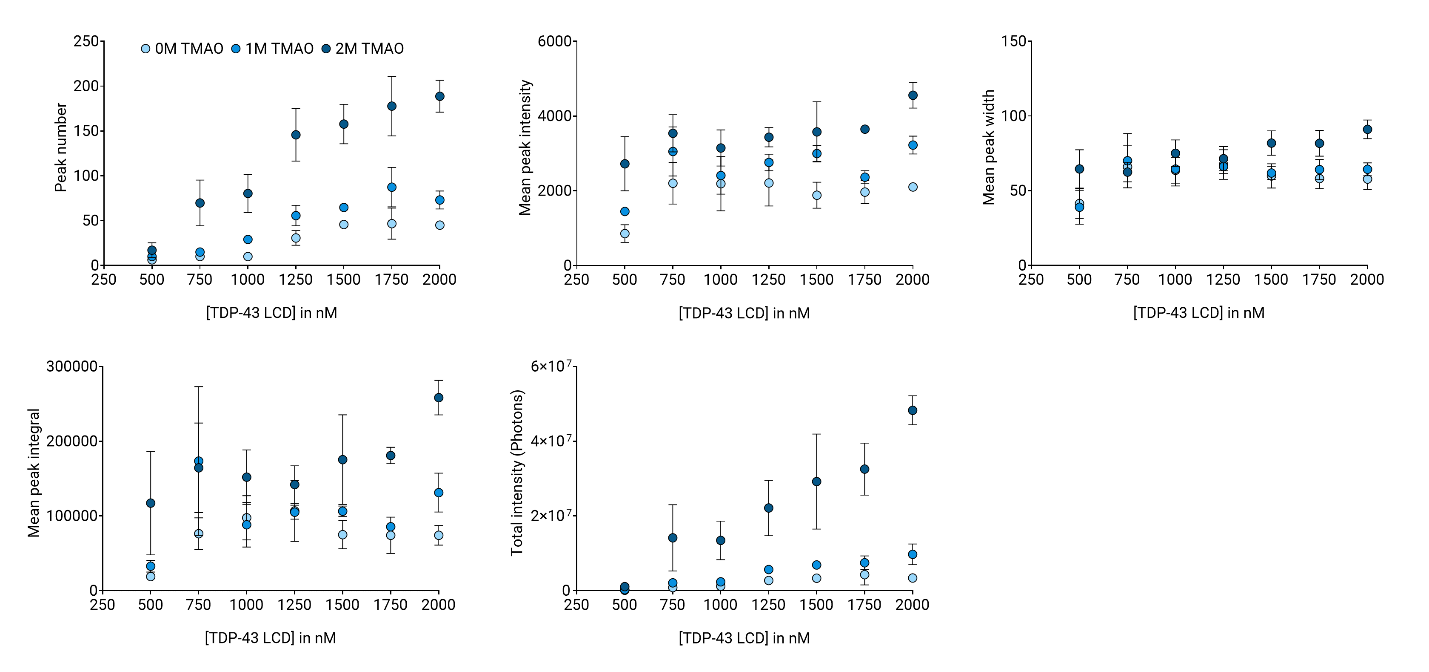

Graphs showing the peak number, mean peak intensity, mean peak width, mean peak integral, and total intensity as a function of TDP-43 LCD concentration, for three separate experiments in buffer with different concentrations of TMAO (0, 1, and 2M in light blue, blue, and dark blue respectively). In all graphs the mean is presented, and error bars are standard error of the mean (SEM).

**Figure S14. Size and internal concentration of condensates formed in 2M TMAO and LTE**

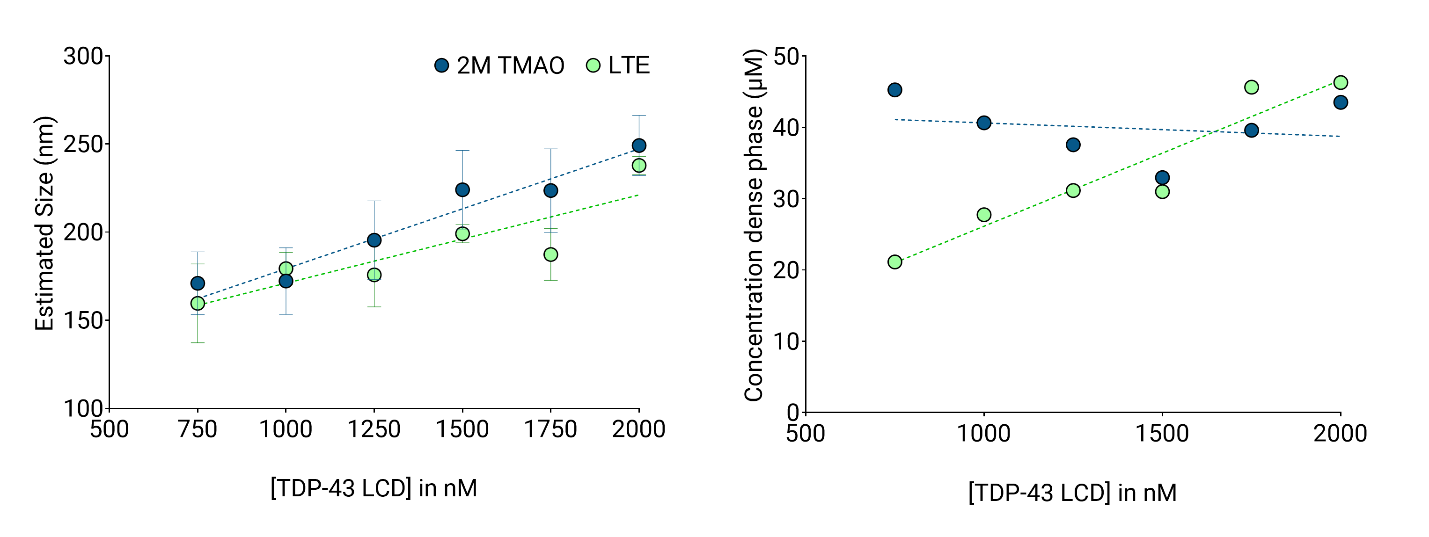

Graphs showing the estimated diameter (based on size calibration) and concentration of TDP-43 LCD inside the condensates formed in either 2M TMAO, αβγ buffer (blue) or cell lysate, LTE (green). In all graphs the mean is presented, and error bars are standard error of the mean (SEM).

**Figure S15. Determination of the volume fraction of protein inside the dense phase**

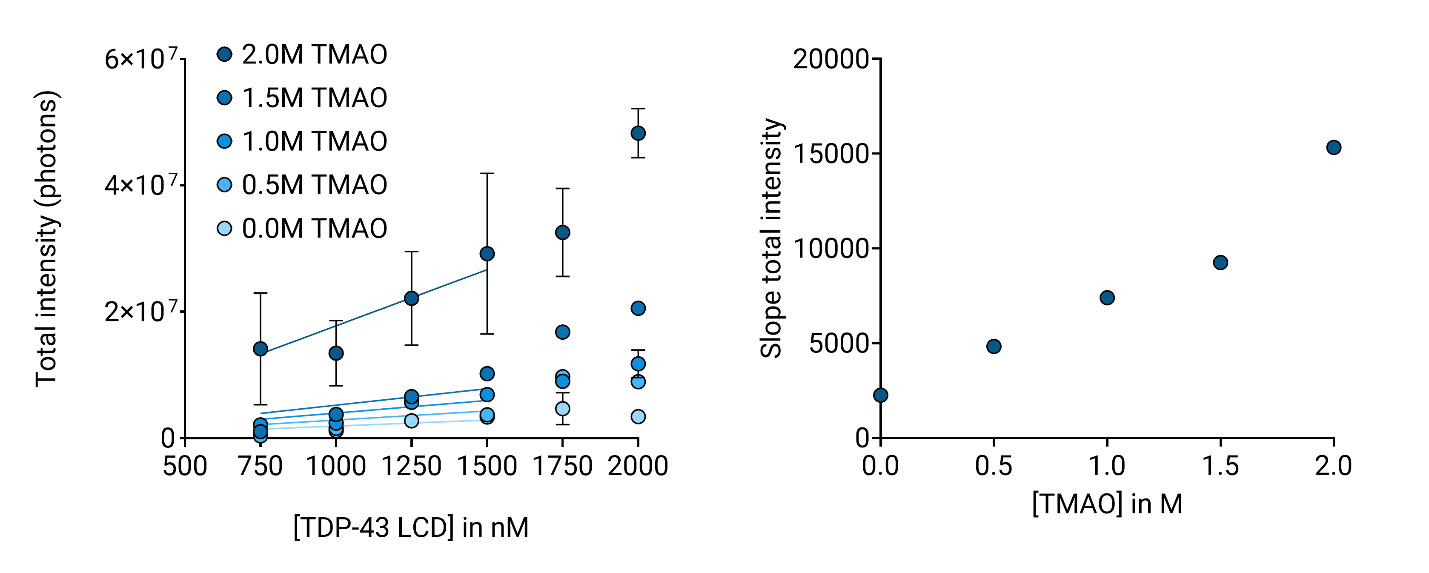

(left) The total intensity in the 10-minute traces varies as a function of total TDP-43 LCD concentration experiments in the presence of different concentrations of TMAO (increasing in concentration from light to dark blue). A linear fit was used to fit the data obtained between 750 and 1500 nM. (right) The slopes obtained for the linear fit in the left figure are plotted as a function of TMAO concentration.

**Figure S16. Evolution of the condensates charcteristics over 1 hour**

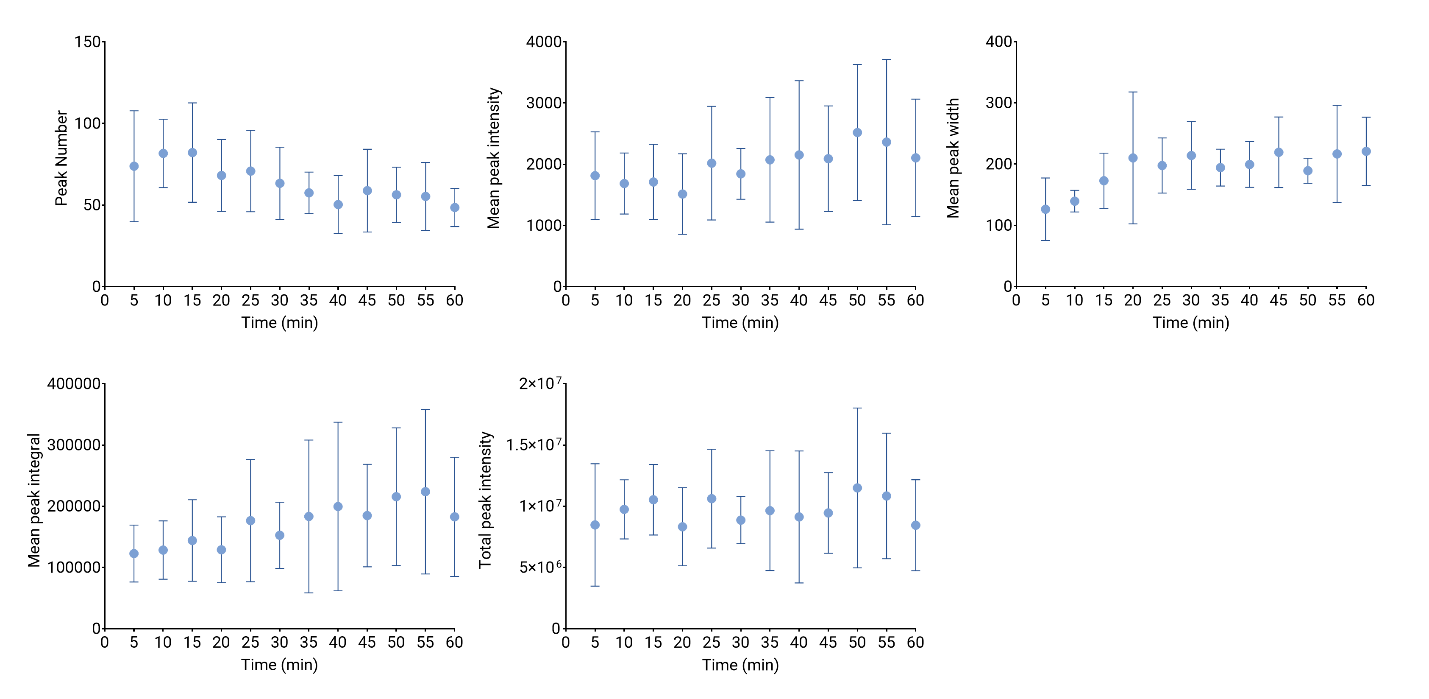

Graphs showing the peak number, mean peak intensity, mean peak width, mean peak integral, and total intensity as a function of time, for six independent 1-hour measurements. The samples contained 500 nM Atto647N-labeled TDP-43 LCD and 500 nM unlabeled TDP-43 LCD in αβγ buffer with 2M TMAO. Data in all graphs show the median and the error bars are standard error of the mean (SEM).

**Table S1. Details of the fluorescent objects used for size calibration**

|  | **Diameter** | **Fluorescence (Ex/Em)** | **Manufacturer / reference** |
| --- | --- | --- | --- |
| HVB capsids purified, labelled with Alexa 488 | 45 nm | 488/525 nm | N/A |
| TetraSpeck™ Microspheres  Ref T7279 | 100 nm | 505/515 nm | Invitrogen |
| Liposomes containing Alexa 488 | 120 nm | 488/ 525 nm | <https://doi.org/10.7554/eLife.74901> |
| FluoSpheres® carboxylate-modified microspheres Ref F8809 | 200 nm | 540/560 nm | Molecular probes |
| SPHERO™ Fluorescent Polystyrene particles Ref FP-0562-2 | 430 nm | 580/620 nm | Spherotech, Inc. |

**SUPPLEMENTARY INFORMATIONS**

**Simulations.**

To improve the understanding of the signals obtained in experiments we performed simulations of diffusion of nanocondensates in the focal volume. In these simulations different parameters can be controlled and their effect on the resulting signal can be studied. In the simulations a spherical shape is considered and the radius r_p_ (radius of particle) can be altered. We assume that the sphere has a free Brownian motion, and its diffusion coefficient can be calculated with the Stokes-Einstein relation:

$$D_{sphere}= \frac{k_{b}T}{6\pi\eta r_{p}}$$

where k_b_ is Boltzmann constant, $T=293K$ is the temperature and $\eta=1.1 \times{10}^{-3}$Pa.s is the dynamic viscosity of the buffer.

To generate a trajectory over time $t$, we divide it in $N$ time steps separated by $dt=t/N$. A tri-dimensional vector with coordinates following a Gaussian distribution of scale $\sqrt{D_{sphere}dt}$ is used for the movement between two time steps. We generate a ($N\times3$) array containing all increments of movement and sum it progressively to obtain a ($N\times3$) array with all the positions. We shift the whole trajectory by a random vector (with a large scale) to have different starting points for each simulation. To simulate several spheres, we constrain the movement of one sphere to a box with periodic boundary conditions. The simulations carried out in this study used a cubic box with a $4 \times4 \times4 \mu m$ dimensions. We assume that the fluorescent markers are homogeneously distributed within the sphere and solution. The intensity observed by the detector should therefore be proportional to the volume of the sphere.

The detector itself has a Gaussian sensitivity with different scales in $z$ and $r$, where $z$ is the axis of the microscope. Therefore, in Cartesian coordinates, it is proportional to:

$$f\left( x,y,z \right)=\exp\left\{ -\frac{z^{2}}{\Delta z^{2}}-\frac{x^{2}+ y^{2}}{\Delta r^{2}} \right\}$$

where $\Delta z=1\mu m$and $\Delta r=0.2\mu m$.

To compute the signal for each position of the sphere, we integrate the sensibility function over the volume of the sphere. The integration is made as a simple Riemann sum, by dividing the sphere in 125,000 boxes and summing the values of the sensitivity function computed at each position. To simulate an acquisition time of 1ms, we generate signals with $dt=$ 0.1ms and we sum the signal by slices of 1ms.

**Calculation of volume and concentrations.**

The mean peak width of different fluorescence traces was converted to mean diameter of the condensates using the previously determined calibration curve using the following equation. Simply by dividing the mean peak width with:

$$D_{sphere}=\omega_{avg}/0.3655$$

where $\omega_{avg}$ is the mean peak width of the peaks in the trace in ms and 0.3655 is the slope of the calibrate curve. Now the volume could be determined from the average size using:

$$V_{sphere}= \frac{4}{3}\pi{r_{sphere}}^{3}$$

where $r_{sphere}$is the average radius of the condensates in the sample in nm.

To calculate the average amount of protein inside the condensates several factors have to be taken into account, which leads to the following equation to determine the average amount of protein in mol:

$$\#mol=\frac{\left( \frac{\left( \frac{{Int}_{avg}}{s_{avg}} \right)}{I_{LAB}} \right)}{N_{A}}\times\frac{\left[ P_{LAB} \right]+\left[ P_{UNL} \right]}{\left[ P_{LAB} \right]}$$

Where $\#mol$ is the average amount of protein in mol, $N_{A}$ is the Avogadro’s number ($N_{A}= 6.02214076\times{10}^{23} {mol}^{-1}$), ${Int}_{avg}$ is the mean peak integral, $\omega_{avg}$ is the mean peak width, $I_{LAB}$ is the intensity of a single fluorophore ($I_{LAB}=52 photon.{ms}^{-1})$, based on PCH (see below), $\left[ P_{LAB} \right]$ is the concentration of labelled protein, and $\left[ P_{UNL} \right]$ the concentration of unlabelled protein both in nM. Finally, the average concentration of protein in the condensates can be determined by using the following formula:

$$\left[ P_{condensate} \right]= \frac{\#mol}{V_{sphere}}$$

where $V_{sphere}$ is the average volume of the condensates in L.

**Photon counting histogram and brightness of Atto647N.**

To calibrate the brightness of the AF647 dye in our setup, a serial dilution of AF647 was performed in our standard buffer (1x αβγ buffer, 0M TMAO) from 250 nM to 1 nM. The samples were measured for 1 minute in 3 separate files of 20 seconds each. The data obtained at the lowest concentrations, where individual fluorophores are detected, where analyzed by photon counting histogram (PCH) analysis. Histograms showing the fraction of events as a function of intensity were obtained and showed that the AF 647 curves converged to a maximal value of approx. 52 photons (Figure S.17).

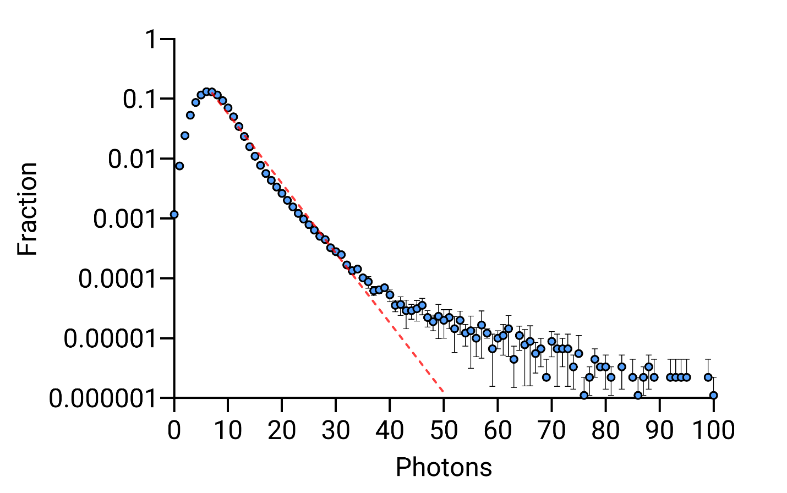
**Figure S17. Photon counting histogram for Atto647N**

Histogram showing the fraction of events as a function of intensity, obtianed from 3 measurements. The red line intersect is a measure of the brightness of the fluorescent dye.

**PYTHON SCRIPTS**

**Quadrant analysis:**

To define 4 sub-quadrants as shown in Figure 5A, thresholds for the peak width and intensity must be determined. Two methods can be used:

1. Reference based fixed values for width and intensity based on a reference trace. A reference trace needs to be obtained, usually in conditions we wanted to use as a comparison for the other parameters of the experiment. For example, in the case of the TDP-43 titration, this would be the lowest concentration of unlabeled TDP-43 LCD or the first 5 minutes of a 1hr measurement. Thresholds are then determined with the requirement that 90% of the events in the reference trace have a peak width and intensity below the defined thresholds.
2. Manually fixed values for width and intensity. In some cases, the reference trace seemed unsuited to determine realistic thresholds. This is the case of the experiments with 0M TMAO where no LLPS occurs. Therefore, it would not make sense to use this condition to determine the intensity and width thresholds. Thus, in the absence of reference, the thresholds are determined manually based on the main spread in the plots for peak intensity and peak width. We expect most peaks at low concentrations to be in the BL quadrant and as time or concentrations increase it is expected the spread of points shifts towards other quadrants. With these criteria in mind thresholds were set at 100ms for width and 2000 counts per ms for intensity.

**Script “quadrant analysis”**

\# -*- coding: utf-8 -*-

"""

@author: jhoux

"""

#%% Import packages

import numpy as np

import pandas as pd

import matplotlib.pyplot as plt

import os

from os import chdir

#%% Retrieve the names of the files in the current directory. We only keep csv files.

c = os.getcwd()

repertoire = os.walk(c)

l = []

for root, dirs, files in repertoire :

l.append(files)

l = l[0]

l = [i for i in l if i.endswith(".csv")]

l = [i for i in l if i != "quadrant_results.csv"]

print(l)

#%%

######## OPTIONS FOR THRESHOLDS AND PLOTS ########

\# Parameters to set the thresholds for the width and the intensity of the peaks.

#Choose if you want to use a manual threshold (If False a reference file is needed named ref.csv)

manual_threshold = True # If True change the following values to the values you want to use as thresholds for the width and Intensity.

w_threshold = 100

i_threshold = 1000

#Choose the proportion of peaks that have to be in a certain quadrant (only used when manual_threshold is turned to False)

width_proportion = .9 # The threshold is set so that at least x% of the peaks in the reference file have a width smaller than the threshold.

intensity_proportion = .9 # The threshold is set so that at least x% of the peaks in the reference file have an intensity smaller than the threshold.

same_scale = True # If True, the graphs will have the same scale for the x and y axes.

save_fig = True # If True, the graphs will be saved in the current directory.

add_text = False # If True, the text will be added to the graphs.

#Choose if you want Bar plots

QA_Bar_Graph = True # If True, a bar graph will be created showing the % of peaks in each quadrant for each file + a table with corresponding filenames.

#Choose if you want plots for both the percentage and number of peaks

perc_bars = True

nr_bars = True

min_width = np.inf

max_width = -np.inf

min_intensity = np.inf

max_intensity = -np.inf

for name in l:

df = pd.read_csv(name)

df.set_index("Unnamed: 0", inplace=True)

df = df.T

widths = df["Peak width"].values

sorted_widths = np.sort(np.copy(widths))

intensities = df["Peak intensity"].values

sorted_intensities = np.sort(np.copy(intensities))

min_width = min(min_width, sorted_widths[0])

max_width = max(max_width, sorted_widths[-1])

min_intensity = min(min_intensity, sorted_intensities[0])

max_intensity = max(max_intensity, sorted_intensities[-1])

if name[:3] == "0.0" or name == "ref.csv": # the reference files is the one starting with 0.0 or the one named ref.csv

n = 0

N = len(widths)

width_threshold = 0

\# The threshold is the width of the first peak wider than 90% of the peaks in the reference file.

while n < width_proportion*N:

width_threshold = sorted_widths[n]

n += 1

intensity_threshold = 0

n = 0

\# The threshold is the intensity of the first peak more intense than 90% of the peaks in the reference file.

while n < intensity_proportion*N:

intensity_threshold = sorted_intensities[n]

n += 1

if manual_threshold:

width_threshold = w_threshold

intensity_threshold = i_threshold

tl_list, tr_list, bl_list, br_list = [], [], [], [] # Top left, top right, bottom left, bottom right lists. Percentages of peaks in each quadrant for each file.

ntl_list, ntr_list, nbl_list, nbr_list = [], [], [], []

for name in l:

tl, tr, bl, br = 0., 0., 0., 0.

df = pd.read_csv(name)

df.set_index("Unnamed: 0", inplace=True)

df = df.T

widths = df["Peak width"].values

intensities = df["Peak intensity"].values

N = len(widths)

for i in range(N):

if intensities[i] >= intensity_threshold:

if widths[i] <= width_threshold:

tl += 1

else:

tr += 1

else:

if widths[i] <= width_threshold:

bl += 1

else:

br += 1

ntl_list.append(tl)

ntr_list.append(tr)

nbl_list.append(bl)

nbr_list.append(br)

tl, tr, bl, br = 100 * tl/N, 100 * tr/N, 100 * bl/N, 100 * br/N # Percentages of peaks in each quadrant.

text = f"Top left : {tl:.2f}%\nTop right : {tr:.2f}%\nBottom left : {bl:.2f}%\nBottom right : {br:.2f}%" # Text to be displayed on the graph

tl_list.append(tl)

tr_list.append(tr)

bl_list.append(bl)

br_list.append(br)

print(tl_list, tr_list, bl_list, br_list)

fig = plt.figure() # Plot the results for each file.

ax = fig.add_subplot(111)

if same_scale:

delta_width = max_width - min_width

delta_intensity = max_intensity - min_intensity

ax.set_xlim(min_width - delta_width/20, max_width + delta_width/20)

ax.set_ylim(min_intensity - delta_intensity/20, max_intensity + delta_intensity/20)

ax.plot(widths, intensities, ".", color="black", markersize=7)

ax.set_xlabel("Peak width", fontsize=12)

ax.set_ylabel("Peak intensity", fontsize=12)

ax.set_title(name[:-4])

ax.tick_params(labelsize=10)

x1, x2 = ax.get_xlim()

y1, y2 = ax.get_ylim()

ax.plot([width_threshold, width_threshold], [y1, y2], "--", color="red", linewidth=1.0)

ax.plot([x1, x2], [intensity_threshold, intensity_threshold], "--", color="red", linewidth=1.0)

if add_text:

ax.text(x1 + .6*(x2 - x1), y1 + .75*(y2 - y1), text, fontsize=16)

if save_fig:

fig.savefig(name[:-4] + ".png", dpi=300)

name_list = [name[:-4] for name in l]

final_dic = {"% Top left" : tl_list, "% Top right" : tr_list, "% Bottom left" : bl_list, "% Bottom right" : br_list, "Npeaks Top left" : ntl_list, "Npeaks Top right" : ntr_list, "Npeaks Bottom left" : nbl_list, "Npeaks Bottom right" : nbr_list}

final_dataframe = pd.DataFrame(final_dic, index=name_list)

final_dataframe.to_csv("quadrant_results.csv", sep=",") # Save the results in a csv file.

\# %%

if QA_Bar_Graph:

if perc_bars:

\# Labels for the x-axis

legend_labels = ['BL', 'BR', 'TL', 'TR']

\# Convert the data to a numpy array

tl = np.array(tl_list)

tr = np.array(tr_list)

bl = np.array(bl_list)

br = np.array(br_list)

\# Number of files

num_files = len(tl)

\# Width of each bar

bar_width = 0.35

\# Position of bars on x-axis

x = np.arange(num_files)

\# Define colors for each quadrant

colors = ['#BADE86', '#567A21', '#A7BEE2', '#324A96']

\# Create a new figure

fig1 = plt.figure(figsize=(10, 10), dpi=300)

\# Adding subplot for the bar graph

plt.subplot(2, 1, 1)

\# Plotting stacked bars with specified colors

plt.bar(x, bl, label='BL', width=bar_width, color=colors[0], edgecolor='black')

plt.bar(x, br, label='BR', width=bar_width, bottom=bl, color=colors[1], edgecolor='black')

plt.bar(x, tl, label='TL', width=bar_width, bottom=bl+br, color=colors[2], edgecolor='black')

plt.bar(x, tr, label='TR', width=bar_width, bottom=bl+br+tl, color=colors[3], edgecolor='black')

\# Adding labels and title

plt.xlabel('File Number')

plt.ylabel('% Peaks')

plt.title('Quadrant Analysis Graph')

plt.xticks(x, range(1, num_files + 1))

\# Displaying the legend on the right of the figure but outside the graph

plt.legend(title='Quadrants', labels=legend_labels, bbox_to_anchor=(1.05, 1), loc='upper left')

\# Adding subplot for the table

plt.subplot(2, 1, 2)

\# Adding a table with filenames and file numbers

table_data = [[i+1, name] for i, name in enumerate(l)]

table = plt.table(cellText=table_data, colLabels=['File Number', 'File Names'], cellLoc='center', loc='center', colWidths=[0.1, 0.5])

\# Adjusting table properties

table.auto_set_font_size(False)

table.set_fontsize(10)

table.scale(2,1)

\# Hide the axes

plt.axis('off')

\# Adjust layout

plt.tight_layout()

\# Show plot

plt.show()

fig1.savefig('Bar_graph_percentage' + ".png", dpi=300)

if nr_bars:

\# Labels for the x-axis

legend_labels = ['BL', 'BR', 'TL', 'TR']

\# Convert the data to a numpy array

tl = np.array(ntl_list)

tr = np.array(ntr_list)

bl = np.array(nbl_list)

br = np.array(nbr_list)

\# Number of files

num_files = len(tl)

\# Width of each bar

bar_width = 0.35

\# Position of bars on x-axis

x = np.arange(num_files)

\# Define colors for each quadrant

colors = ['#BADE86', '#567A21', '#A7BEE2', '#324A96']

\# Create a new figure

fig2 = plt.figure(figsize=(10, 10), dpi=300)

\# Adding subplot for the bar graph

plt.subplot(2, 1, 1)

\# Plotting stacked bars with specified colors

plt.bar(x, bl, label='BL', width=bar_width, color=colors[0], edgecolor='black')

plt.bar(x, br, label='BR', width=bar_width, bottom=bl, color=colors[1], edgecolor='black')

plt.bar(x, tl, label='TL', width=bar_width, bottom=bl+br, color=colors[2], edgecolor='black')

plt.bar(x, tr, label='TR', width=bar_width, bottom=bl+br+tl, color=colors[3], edgecolor='black')

\# Adding labels and title

plt.xlabel('File Number')

plt.ylabel('Number of peaks')

plt.title('Quadrant Analysis Graph')

plt.xticks(x, range(1, num_files + 1))

\# Displaying the legend on the right of the figure but outside the graph

plt.legend(title='Quadrants', labels=legend_labels, bbox_to_anchor=(1.05, 1), loc='upper left')

\# Adding subplot for the table

plt.subplot(2, 1, 2)

\# Adding a table with filenames and file numbers

table_data = [[i+1, name] for i, name in enumerate(l)]

table = plt.table(cellText=table_data, colLabels=['File Number', 'File Names'], cellLoc='center', loc='center', colWidths=[0.1, 0.5])

\# Adjusting table properties

table.auto_set_font_size(False)

table.set_fontsize(10)

table.scale(2,1)

\# Hide the axes

plt.axis('off')

\# Adjust layout

plt.tight_layout()

\# Show plot

plt.show()

fig2.savefig('Bar_graph_number' + ".png", dpi=300)

**Script: “first peak analysis”-** for data analysis as in Figure 4

\# -*- coding: utf-8 -*-

"""

@author: jhoux

"""

import os

import pandas as pd

\# Function to determine the first peak position above a given intensity threshold

def first_peak_position_above_intensity(intensity_threshold, peak_intensities, peak_positions):

for intensity, position in zip(peak_intensities, peak_positions):

if intensity > intensity_threshold:

return position

return None # Default peak position if no peak is found

\# Function to process each raw file

def process_raw_files(input_folder, intensity_values):

results = {}

for filename in os.listdir(input_folder):

if filename.endswith(".csv") and filename != 'results.csv':

filepath = os.path.join(input_folder, filename)

df = pd.read_csv(filepath, index_col=0)

peak_positions = df.loc["Peak position"].values

peak_intensities = df.loc["Peak intensity"].values

file_results = []

for intensity in intensity_values:

peak_position = first_peak_position_above_intensity(intensity, peak_intensities, peak_positions)

file_results.append(peak_position)

results[filename] = file_results

return pd.DataFrame(results, index=intensity_values)

\# Get the current directory

current_directory = os.getcwd()

\# Define the range of intensity values

max_intensity = 20001

step_size = 500

intensity_values = range(0, max_intensity, step_size)

\# Process files in the current directory

results_df = process_raw_files(current_directory, intensity_values)

\# Define the output file path

output_file = os.path.join(current_directory, "results.csv")

\# Save results to the output file

results_df.to_csv(output_file)

**Script: “single molecule analysis”**- to retrieve peak number, intensity, width and area under the curve

\# -*- coding: utf-8 -*-

"""

@author: ygamb

"""

###########################################################################

\## Librairy

###########################################################################

import matplotlib.pyplot as plt

import numpy as np

import matplotlib.image as mpimg

import os

from skimage import exposure

from scipy.ndimage import gaussian_filter

from skimage.morphology import reconstruction

import pandas as pd

\# import cv2

from skimage import color

from os import chdir

import scipy.signal as signal

from scipy.signal import butter,filtfilt

from matplotlib.colors import LogNorm

import re

###########################################################################

\## Recuperation des fichiers

###########################################################################

c = os.getcwd()

repertoire = os.walk(c)

liste_elem = []

for root, dirs, files in repertoire :

liste_elem.append([root, files])

###########################################################################

\## Traitement des donnees

###########################################################################

def butter_lowpass_filter(data, cutoff, fs, order):

normal_cutoff = cutoff / nyq

\# Get the filter coefficients

b, a = butter(order, normal_cutoff, btype='low', analog=False)

y = filtfilt(b, a, data)

return y

###########################################################################

\## reccuperer les pics pics

###########################################################################

##Step 1 : Define the filter requirements

\# Filter requirements.

T = 12000	# Sample Period

fs = 1000.0	# sample rate, Hz

cutoff = 20	# desired cutoff frequency of the filter, Hz , slightly higher than actual 1.2 Hz

nyq = 0.5 * fs # Nyquist Frequency

order = 2	# sin wave can be approx represented as quadratic

n = int(T * fs) # total number of samples

rapport = 20

ref1 = 0

taille = 200 # Calcul ecart type

mean_intensity = []

mean_width = []

mean_intengral = []

liste_nbr_peak = []

liste_name = []

liste_rapp = []

liste_mean_fin = []

#Step 2 : Filter implementation using scipy

for elem in liste_elem :

chdir(elem[0])

liste_fichier = elem[1]

for fichier in liste_fichier :

if fichier[len(fichier)-5:] != ".xlsx" and fichier[len(fichier)-4:] != ".png" and fichier[len(fichier)-4:] != ".csv" and fichier[len(fichier)-3:] != ".py" and fichier[len(fichier)-4:] != ".txt":

liste = pd.read_table(fichier)

label = liste.columns[0]

liste_y = liste[label].tolist()

y = butter_lowpass_filter(liste_y, cutoff, fs, order)

liste_x = np.linspace(0,len(liste_y)-1,len(liste_y))

liste_prom = []

liste_mean = []

for i in range (1,len(y)-taille,taille):

liste_prom.append(np.std(liste_y[i:i + taille])*rapport)

liste_mean.append(np.mean(liste_y[i:i + taille]))

prominence_min = min(liste_prom)

back = liste_mean[liste_prom.index(prominence_min)]

liste_mean_fin.append(back)

bruit = back/(prominence_min/rapport)**2

peaks, properties = signal.find_peaks(y, prominence =(prominence_min,None))

widths = signal.peak_widths(y, peaks)

liste_p = []

for p in peaks :

liste_p.append(y[p])

liste_left = widths[2]

liste_right = widths[3]

liste_intensite = []

liste_integrale = []

for i in range(len(peaks)) :

left = int(liste_left[i])

right = int(liste_right[i])

liste_integrale.append(sum(liste_y[left:right]))

liste_intensite.append(max(liste_y[left:right]))

fig = plt.figure(figsize=(50,30))#, dpi=300)

plt.plot(liste_x,liste_y, label = "prominence : " + str(prominence_min)) # Orange Line --> proccessed trace

plt.plot(liste_x,y, label = "background : " + str(back)) # blue line --> raw trace

#plt.ylim(0,20000)

plt.legend(fontsize = 60)

plt.ylabel("Intensity (Photons/ms)", fontsize = 60)

plt.xlabel("Time (ms)", fontsize = 60)

plt.title(fichier, fontsize = 80)

plt.tick_params(labelsize = 60)

#x_length = len(liste_x) + 1

#x_minutes = int(x_length/60000)

#x_labels = list(range(x_minutes + 1))

#plt.xticks(np.arange(0, len(liste_x) + 2, 60000), x_labels)

#plt.tick_params(axis='both', which='both', length=20, width=3) # Set thickness for ticks on both axes

#plt.tick_params(axis='both', which='minor', length=10, width=2) # Set thickness for minor ticks

#plt.gca().spines['top'].set_linewidth(2) # Set thickness for top spine

#plt.gca().spines['right'].set_linewidth(2) # Set thickness for right spine

#plt.gca().spines['bottom'].set_linewidth(2) # Set thickness for bottom spine

#plt.gca().spines['left'].set_linewidth(2) # Set thickness for left spine

if bruit >= ref1 :

plt.plot(peaks, liste_p, 'o', markeredgecolor = "r", markersize = 20, label = "rapport : " + str(bruit))

plt.legend(fontsize = 60)

data = [[len(peaks)],

peaks,

liste_intensite,

widths[0],

liste_integrale,

[back],

[bruit]]

df = pd.DataFrame(data, index = ['Peak number',

'Peak position',

'Peak intensity',

'Peak width',

'Peak intengrale',

'Background',

'Rapport'])

if len(peaks)>0 :

mean_intensity.append(sum(liste_intensite)/len(peaks))

mean_width.append(sum(widths[0])/len(peaks))

mean_intengral.append(sum(liste_integrale)/len(peaks))

liste_nbr_peak.append(len(peaks))

else :

mean_intensity.append(None)

mean_width.append(None)

mean_intengral.append(None)

liste_nbr_peak.append(len(peaks))

else :

plt.axhline(np.mean(liste_y), linestyle= '--', color = 'r', label = 'Mean ='+ str(np.mean(liste_y)))

plt.legend(fontsize = 60)

data = [[np.mean(liste_y)], [prominence_min/rapport], [sum(liste_y)], [bruit]]

df = pd.DataFrame(data, index = ['Mean',

'Std',

'Integral',

'Rapport'])

mean_intensity.append(np.mean(liste_y))

mean_width.append(None)

mean_intengral.append(sum(liste_y))

liste_nbr_peak.append("High concentration")

liste_rapp.append(bruit)

liste_name.append(fichier)

df.to_csv(fichier+".csv")

plt.savefig(fichier+".png")

plt.show()

\# plt.hist(liste_integrale, 20)

\# plt.title("Integrale : " + fichier)

\# plt.savefig(fichier+"_hist integrale.png")

\# plt.show()

\# plt.hist(liste_intensite, 20)

\# plt.title("Intensitee : "+ fichier)

\# plt.savefig(fichier+"_hist intensite.png")

\# plt.show()

liste_itot = []

for i in range(len(liste_nbr_peak)):

itot = liste_nbr_peak[i] * mean_intengral[i]

liste_itot.append(itot)

chdir(liste_elem[0][0])

data = {'Name' : liste_name,

'Peak number' : liste_nbr_peak,

'Mean peak intensity' : mean_intensity,

'Mean peak width' : mean_width,

'Mean peak intengrale' : mean_intengral,

'background' : liste_mean_fin,

'Rapport' : liste_rapp,

'Total peak intensity' : liste_itot}

df = pd.DataFrame(data)

df.to_csv("final_analyse.csv")

liste_fich = np.linspace(0,len(liste_rapp)-1,len(liste_rapp))

fig = plt.figure(figsize=(50,30))

plt.plot(liste_fich,liste_rapp, 'o', markeredgecolor = "r", markersize = 30)

plt.axhline(ref1, linestyle= '--', color = 'b', label = 'rapport ='+ str(ref1))

plt.legend(fontsize = 60)

plt.ylabel("Rapport", fontsize = 60)

plt.xlabel("Files", fontsize = 60)

plt.title("Rapport", fontsize = 80)

plt.tick_params(labelsize = 40)

plt.savefig("Rapport.png")

plt.show()

fig = plt.figure(figsize=(50,30))

plt.plot(liste_fich,liste_mean_fin, 'o', markeredgecolor = "r", markersize = 30)

plt.legend(fontsize = 60)

plt.ylabel("Mean", fontsize = 60)

plt.xlabel("Files", fontsize = 60)

plt.title("Mean", fontsize = 80)

plt.yscale("log")

plt.tick_params(labelsize = 40)

plt.savefig("Mean.png")

plt.show()

\# ###########################################################################

\# ## Sauvegarder la taille des images dans un fichier excel

\# ###########################################################################

\# L = blobs_dog

\# dfObj = pd.DataFrame(L)

\# dfObj.to_csv("fichier_ref.csv")
